# Cas9 RNP Physiochemical Analysis for Enhanced CRISPR-AuNP Assembly and Function

**DOI:** 10.1101/2024.04.02.586657

**Authors:** Daniel D. Lane, Karthikeya S.V. Gottimukkala, Rachel A. Cunningham, Shirley Jwa, Molly E. Cassidy, Jack M.P. Castelli, Jennifer E. Adair

## Abstract

CRISPR therapy for hematological disease has proven effective for transplant dependent beta thalassemia and sickle cell anemia, with additional disease targets in sight. The success of these therapies relies on high rates of CRISPR-induced double strand DNA breaks in hematopoietic stem and progenitor cells (HSPC). To achieve these levels, CRISPR complexes are typically delivered by electroporation ex vivo which is toxic to HSPCs. HSPCs are then cultured in stimulating conditions that promote error-prone DNA repair, requiring conditioning with chemotherapy to facilitate engraftment after reinfusion. In vivo delivery by nanocarriers of CRISPR gene editing tools has the potential to mitigate this complexity and toxicity and make this revolutionary therapy globally available. To achieve in vivo delivery, the inherent restriction factors against oligonucleotide delivery into HSPCs, that make ex vivo manipulation including electroporation and stimulation essential, must be overcome. To this end, our group developed a CRISPR carrying gold nanoparticle (CRISPR-AuNP) capable of delivering either Cas9 or Cas12a CRISPRs as ribonucleoprotein complexes (RNP) without compromising HSPC fitness. However, the most commonly used CRISPR, Cas9, demonstrated inconsistent activity in this delivery system, with lower activity relative to Cas12a. Investigation of Cas9 RNP biophysics relative to Cas12a revealed duplex RNA instability during the initial loading onto Au cores, resulting in undetectable Cas9 loading to the particle surface. Here we demonstrate preformation of RNP before loading, coupled with optimization of the loading chemistry and conditions, resulted in 39.6 ± 7.0 Cas9 RNP/AuNP without compromising RNP activity in both in vitro assays and primary human HSPC. The same alterations improved Cas12a RNP/AuNP loading 10-fold over previously reported levels. To achieve particle stability, the reported polyethyleneimine outer coating was altered to include PEGylation and the resulting 2^nd^ generation CRISPR-AuNP demonstrates favorable nanoformulation characteristics for in vivo administration, with a hydrophilic, more neutral nanoparticle surface. Direct treatment of HSPC in vitro showed 72.5 ± 7.37% uptake of 2^nd^ generation CRISPR-AuNP in primary human HSPC, but with endosomal accumulation and low rates of gene editing consistent with low levels of endosomal escape.

## Introduction

Providing durable therapy for blood disorders requires direct genetic manipulation of hematopoietic stem and progenitor cells (HSPCs), the lifelong cell source for blood development.^1^ Therapeutic CRISPR gene editing of HSPCs recently passed a milestone with approval of Casgevy in the U.K. and U.S., using ex vivo manipulated HSPCs to treat sickle cell anemia.^2,3^ However, current state of the art for these therapies requires mobilization and purification of HSPCs combined with multi day cellular manipulation, including stem cell supportive culture, electroporation, and/or engineered lentiviral particle transduction to deliver CRISPR. These processes warrant an International Standards Organization (ISO) class 5 clean room and require patient conditioning prior to reinfusion of these heavily manipulated cell products. While this approach is effective, to date it has not demonstrated feasibility for global dissemination. Facilitating efficient delivery of CRISPR gene editing to HSPC in vivo would increase access, but this ability remains the grail of genetic editing.

In vivo, HSPCs are biologically rare, representing only 0.8% of total nucleated marrow cells.^4^ Direct targeting in peripheral blood, outside the niche, requires either proliferation and spillover into circulation by filgrastim (recombinant human granulocyte colony stimulating factor (G-CSF)) and/or mobilization by plerixafor (AMD3100), which increases the peripheral blood CD34^+^ cell level to 0.2% of nucleated cells.^5,6^ Without conditioning, HSPC homeostasis is maintained through self-renewal with signal directed activation and differentiation for leuko- and myelogenesis.^7^ This required long term pluripotency is theorized to have selected for high resistance to genetic alteration and viral transfection, leading to strong restriction of genetic editing.^8^ In addition, HSPC have low translational activity and high chromosomal accessibility, resulting in little mRNA to protein synthesis, and increased likelihood of genomic off target access.^8,9^ Considering these biological characteristics, the barriers to in vivo HSPC gene editing are stark, even with the most accurate and effective editing tools.

Of the discovered gene editing tools, the reprogrammable CRISPR/Cas enzyme systems have become the most prevalent since Cas9’s description in 2012 by Charpentier and Doudna.^10^ The CRISPR system is a combination of a protein nuclease and guide RNA complex (gRNA), forming an active ribonucleoprotein (RNP). Cas9 was the first described CRISPR system, utilizing a partially complementary dual RNA guide of crispr RNA (crRNA) and trans-activating crispr RNA (trRNA), which form an active duplex (duRNA) within the RNP. Its primary nuclease activity generates blunt-end double-stranded DNA breaks which are highly effective in indel formation. In addition, Cas9 has undergone considerable protein engineering and fusions, producing next generation CRISPR tools including prime editors, base editors, nickases and more.^11–23^ An alternative well-described CRISPR system is Cas12a, which utilizes single gRNA (sgRNA) to target DNA and generates a 5 bp overhang, staggered-end cut. Lower editing efficiency has been reported for Cas12a, but its improved thermal stability, short-length native single guide, and specificity over Cas9 offer considerable advantages.^24–27^

The most notable in vivo delivery methods for CRISPR primarily rely on oligonucleotide mRNA/plasmid gene packaging. The first delivery agents were lentivirus, adeno-associated virus and adeno virus which each have shown initial promise but have failed to reach curative in vivo editing of HSPC in clinical trials.^28–30^ Lipid nanoparticles have gained considerable interest due to COVID vaccines success, being able to deliver oligonucleotides into circulation with strong uptake in liver and muscle cells, and are currently under clinical evaluation.^31^ However, as both viral and lipid nanoparticle systems rely on oligonucleotide delivery, they suffer directly from low translational activity of in situ HSPCs, requiring targeting, large doses, and optimization of the therapeutic window across patients.^32^ Direct improvements to each of these are desired for durable, clinical therapeutic editing.

Combining the limitations of HSPC biology and the need for systemic delivery, we developed a synthetic delivery vehicle able to carry CRISPR/Cas RNP as well as DNA and RNA on a colloidal, spherical gold nanoparticle (AuNP).^33^ RNP were anchored through a thiol modification of du/gRNA which can form a self-assembled monolayer on the Au particle surface. After RNA monolayer formation, Cas9/12a nuclease was added and allowed to associate, forming tethered RNP on the Au surface. Finally, these particles were coated in polyethylenimine (PEI), an endosomal disrupting polymer with net positive charge, which permitted loading of additional DNA.^34^ This CRISPR-AuNP passively entered and produced detectable gene editing at two different loci, the *γ*-globin (*HBG1/2*) promoter and the C-C chemokine receptor type 5 (*CCR5*) delta32 mutation site, in primary human HSPCs in vitro with non-statistical difference in toxicity over untreated cells. However, levels of gene editing observed with Cas9 was on par with Cas12a, which is known to have a lower genetic editing rate. Therefore, we sought to investigate Cas9’s physiochemistry and interaction with the surface of the gold to improve CRISPR-AuNP activity.

## Methods

### Materials

All nucleic acids were purchased through IDT (Coralville, Iowa, US). NLS-SpCas9-NLS and NLS-AsCas12a-NLS were purchased from Aldevron (Fargo, North Dakota, US). The chemicals chloroauric acid, sodium citrate dibasic trihydrate, >37% hydrochloric acid, 50% branched 2k PEI, and β-mercaptoethanol (BME) were purchased from Sigma-Aldrich (St. Louis, Missouri, US). For electroporations, the Neon™ Electroporation System 10-μL Kit (Invitrogen, Waltham, MA, USA) was used on the Neon Transfection System (Invitrogen). Electroporation kit V and Electroporation kits for Primary T-Cell/HSPC were purchased from Lonza (Basel, Switzerland) and used on the Lonza Nucleofector 2b system. All oligonucleotides were purchased from Integrated DNA Technologies (Coralville, Iowa, USA). All antibodies were purchased from Biolegend (San Diego, California, US). Other materials not listed were purchased through Thermo-Fischer Scientific (Waltham, Massachusetts, US).

### Au core Synthesis

Spherical Au nanocores were generated via Turkevich’s method.^35,36^ Briefly, chloroauric acid solution (0.25 mM) was brought to its boiling point and then reduced with 3.33% sodium citrate solution under rapid mixing for 15 min. After the solution developed a ruby color, batches were removed from heat and allowed to reach room temperature overnight. The hard diameter of Au cores was determined by transmission electron microscopy (TEM; Talos L120C, ThermoFisher Scientific). Concentration was determined with 1 cm Beer–Lambert law absorbance at 520 nm (Nanodrop One, Thermo-Fischer) with published Au core extinction coefficients.^37^ Batches of Au cores were then stored at 4°C without further refinement.

### 1st generation CRISPR-AuNP Synthesis

CRISPR/AuNP were generated according to the method published by Shahbazi et. al.^38^ Au cores were added to an acid-treated 2mL microcentrifuge tube (Corning, Corning, NY, USA). All incubation steps were carried out at room temperature on the benchtop. All washes utilized an Eppendorf 5702 RH centrifuge at 15 krcf, 15 min at 4°C (Eppendorf, Hamburg, Germany). duRNA was generated through equimolar annealing of trRNA/crRNA in duplex buffer (IDT), [duRNA]_f_ = 50 µM. This duRNA was then added via rapid pipetting to Au cores at 2:1 duplex/Au core mass ratio and mixed well. Citrate buffer (500 mM, pH 3.0) was added to 25 mM and the mixture incubated for 30 min at room temperature. samples were washed with UltraPure Water (UPW; Invitrogen, USA) twice to generate the product designated “AuRNA” stage. A 1.2 μM solution of Cas9 nuclease protein in UPW was generated in a second acid-treated vial to which AuRNA was added and incubated for 10 min, yielding the product designated “AuRNP” stage. 0.1% PEI solution (in UPW, pH 7.0) was added at a 2:1 PEI:Au core mass ratio and quickly mixed via trituration and vortexing before incubating for 10 min and designated “CRISPR-AuNP”. Particles were stored at 4 °C.

### 2nd Generation CRISPR-AuNP Synthesis

2^nd^ Generation CRISPR-AuNP were synthesized by first preforming Cas9/Cas12a RNP. For Cas9 an equimolar solution of cr/trRNA was made in duplex buffer as above, then 2.5:1 duRNA:Cas9 molar solution was made in Dulbecco’s phosphate buffered saline (DPBS; Gibco) to a final RNP concentration of 9.2 μM. For Cas12a, a 2.5:1 gRNA:Cas12a molar solution was made as described for Cas9. These RNP solutions were incubated overnight at 4 °C before adding to Au cores at various Au core/RNP mass ratios as described. The solution was incubated on a shaker for 1 hr at room temperature. A 500 mM, pH 3.8 citrate solution was added to the AuRNP solution to a final concentration of 25 μM. This solution was incubated for 5 min before being washed at 5 krcf for 45 min at 4 °C. The product was resuspended into pH 7.4, 10 mM HEPES buffer and designated “AuRNP” stage. A solution of branched PEI 2k MW-g 10%-PEG 2kSH (PPSH; Nanosoft Polymers, Winston-Salem, NC, USA) was made to 1 mg/mL in HEPES buffer, added to AuRNP before incubating over 15 min. Particles were stored at 4°C.

### Dynamic Light Scattering (DLS) and Transmission Electron Microscopy (TEM)

Particle size and zeta potential were quantified with a Malvern Zetasizer Nano-ZS (Malvern, Worcestershire, UK) at 10 μg/mL in formulation buffer (UPW/HEPES) for DLS, or 10 mM HEPES for zeta potential. DLS readings were three averaged runs of 10 sub runs at 25 °C in BRAND 40 μL disposable cuvettes (BrandTech Scientific, USA). Zeta potential measurements were averaged over 3 reads at 20°C using autosense for each read in bent capillary cells (Malvern). Particle TEM was run on Talos L120C using 1% uranyl acetate as the negative stain. Au core diameter was quantified by drawing a vertical and horizontal cross over each particle and averaging these measurements (n = >10 particles).

### DNA Cutting Assay

DNA cutting assays were performed following New England Biolabs (NEB) method.^39,40^ Briefly, genomic DNA was extracted from primary isolated CD34^+^ cells (STEMCELL Technologies, Vancouver, BC, Canada) using a PureLink® Genomic DNA Minikit (Invitrogen). The target amplicon DNA containing the desired CRISPR cut site was extracted using primers (Supplement Table 1), amplified with Q5® Hot Start 2x Master Mix (New England Biolabs, Ipswich, Massachusetts, US). Target DNA was purified with a PureLink® PCR cleanup kit (Invitrogen). 500 ng of target DNA was then added to PCR strip tubes, along with 1x NEB 3.1r buffer (New England Biolabs). Cas9/Cas12a preformed RNP was a positive control and AuNP samples at various stages were added at 5 ug Au core ± BME (Sigma-Aldrich), V_t_ = 50 μL. Samples were vortexed before incubation at 37 °C for 15 min. Samples then had 1 μL Proteinase K and 1 μL RNase A added (both from Invitrogen) and were incubated for 10 min at 28 °C, then 10 min at 32 °C before being heated to 95 °C to denature and cooled to 23 °C over 15 min. Resulting DNA samples were resolved for size on TapeStation (Agilent, Santa Clara, California, US) for quantification and 2% agarose gel electrophoresis for visualization.

### Kinetic Fluorescence Recovery of trRNA/sgRNA

1^st^ Generation AuRNA were generated with either gRNA-ATTO550 for Cas12a AuNP, or trRNA-ATTO550 for Cas9 AuNP. These particles were suspended in UPW at 25 μg/mL Au core, then placed into black chimney wall, clear flat bottom polystyrene 96 well plates (Greier Bio One, Kremsmünster, Austria). The baseline fluorescence of each well was read for 10 min using a Spark 10M Fluorescent Plate Reader (Tecan, Männedorf, Switzerland), excitation/emission were 550/595, gain 100, 20 nm slit. BME (C_f_=55 μM) or UPW was then added to a total volume of 250 μL and the fluorescence tracked over 16 hours by kinetic loop.

### Fluorescent tr/crRNA Loading Quantification

The level of cr/trRNA binding were simultaneously determined by generating 1^st^ generation AuRNA using crRNA-ATTO488 and trRNA-ATTO550. For all RNA-only controls, Au core storage solution was used after removing the gold cores at 21 krcf for 10 min at 4 °C to mimic synthesis conditions. Following each centrifugation supernatant was reserved. The final AuRNA samples had bound RNA released by BME at 100 μM with overnight incubation at 4 °C. After BME release, Au cores were spun out at 21 krcf, 10 min at 4 °C to remove interference. Supernatants were then diluted into pH 7.4 DPBS and all trRNA-ATTO550 in the supernatant quantified using the Spark 10M plate reader.

### SDS-PAGE Cas Nuclease Binding Determination

CRISPR-AuNP were synthesized as previously described. After synthesis, the nanoformulation was pelleted (5 krcf, 45 minutes, 4°C) and supernatant removed. Particles were then resuspended in 5 mM BME in DPBS for overnight incubation at 4°C to release RNP from the Au core. Au cores were removed from the supernatant by centrifugation, then SDS-PAGE was performed using NuPAGE™ 4-12% Bis-Tris Mini Protein Gels (Invitrogen) according to the manufacturer’s protocol. Gels were stained with SimplyBlue SafeStain (LC6060) using the manufacturer’s protocol and imaged using an Invitrogen iBright Imaging System. Band intensity was quantified using ImageJ software (version: 1.51j8) and compared to a standard curve generated with known amounts of Cas9 nuclease.

### Cas9 pH Stability Quantification

Cas9 and Cas12a preformed RNPs were formulated 1 day before the assay. Preformed Cas9 RNP was prepared in Protein LoBind tubes (Eppendorf) according to the 2^nd^ generation CRISPR-AuNP synthesis protocol.

500 µL of Au cores (45 mg/mL) and 6.7 µL of Cas9 RNP or 4.3 µL of Cas12a RNP were mixed in Protein LoBind tubes and placed on a table shaker at low-medium speed for 1 hour at room temperature. 500 mM citrate buffer was formulated in 2 mL Eppendorf tubes by adding citrate to UPW and HCl (>37%) to achieve the final pH of 2.5-4 for Cas9 and 2-3.5 for Cas12a. Citrate was combined to 25 mM with AuNP solution and samples were incubated for 15 minutes at room temperature. Samples were spun at 5 krcf for 45 minutes at 4 °C, and supernatant was removed. Pellets were resuspended in 100 µL of 5 mM BME in 1x DPBS, then incubated on an orbital shaker at 300 rpm for 1 hour at 37 °C. Au cores were then removed by centrifugation at 5 krcf for 15 minutes at 4 °C. 70 µL of supernatant was collected in Protein LoBind tubes for DNA cutting assay as described above.

### β*-2 microglobulin* (*B2M*) gRNA Optimization

All possible *B2M* exonic gRNA were identified using Benchling [Biology Software] (Version: 2023-24, San Francisco, California, USA). From these guides, 20 were selected using combined Lindel Cas9 prediction and homology directed repair likelihood based on Tatiossian et al.^23,41^ An additional 6 gRNA were pulled from the protomer region which Lindel did not identify.^42^

Jurkat cells (E6-1, ATCC, Manassas, Virginia, USA) were cultured in Gibco Roswell Park Memorial Institute (RPMI)-1640 media + 10% fetal bovine serum (FBS) + 1% Penicillin/Streptomycin (ThermoFisher) at 5% CO_2_, 37 °C, normoxic conditions for 2 passages before use. crRNA was acquired from IDT and Jurkat cells were electroporated according to Lonza’s Cas9 method on the 2b system (120 pmol RNP) into 500k cells/electroporation. Cells were cultured for 3 days, then analyzed by flow cytometry for cell surface B2M protein expression with phycoerythrin (PE)-conjugated anti-human B2M antibody, clone 2M2 (Biolegend). Flow cytometry was performed on a FACSCelesta (BD Life Sciences, Franklin Lakes, NJ, USA) and data were analyzed using FlowJo™ v 10.9 Software (BD Life Sciences).

For DNA sequencing, gDNA was extracted using full *B2M* locus primers (Supplement Table 1) and amplified with Q5® Hot Start 2x Master Mix (New England Biolabs, Ipswich, Massachusetts, US). The target DNA was purified by PureLink® PCR cleanup (Invitrogen), then the target exon was extracted with *B2M* exon primers with attached Nextera XT DNA Library Preparation Kit (Illumina, San Diego, Ca, USA) index adapter sequences. The PCR amplicons were cleaned up and unique indexes attached to the samples before being run on a MiSeq v2 chip (Illumina), read depth >10000 per sample. Genetic editing analysis was completed using either CRISPRESSO or an in-house pipeline previously described.^33,43^

### CD34^+^ Cell Culture

All human CD34^+^ cells were sourced from the Core Center of Excellence in Hematology at the Fred Hutch after mobilization with G-CSF (Amgen, Thousand Oaks, California, USA). Sourcing followed the protocol approved by the Fred Hutch Institutional Review Board (protocol no. 985.03) and in accordance with the Declaration of Helsinki and the Belmont Report. Cells were cultured in StemSpan™ Serum-Free Expansion Medium version II (SFEM II; STEMCELL Technologies) containing 50 ng/mL each of recombinant human stem cell factor (SCF; Peprotech, Cranbury, NJ, USA), thrombopoetin (TPO) and Flt-3 ligand (both from CellGenix, Freiburg, Germany). An overnight pre-treatment incubation was carried out after thaw at 37 °C, 5% CO_2_, and normoxia.

### Direct CRISPR-AuNP 2^nd^ Generation Cell Treatment

Primary human CD34+ HSPCs were thawed from cryopreservation into SFEM II and supplemented with growth factors SCF, Flt-3 ligand and TPO before being pre-stimulated overnight at 37°C in a humidified 5% CO_2_ incubator. 250,000 cells were then seeded at a density of 500,000 cells/mL in 48-well plates and treated with CRISPR-AuNP nanoformulation, controlling for RNP dose. All treatments were performed in triplicate against three independent human cell donors (biological replicates). After 72 hours of incubation at 37°C in a humidified 5% CO_2_ incubator, cells were washed with DPBS and harvested for flow cytometry and genomic DNA (gDNA) extraction for gene editing analysis.

### Electroporation of 2^nd^ Generation CRISPR-AuRNP Released RNP

Second-generation CRISPR-AuNPs were synthesized to the AuRNP stage. The nanoformulation was washed twice in 10 mM HEPES buffer at 5 krcf for 45 min at 4°C, then resuspended in 50 μL of 10 mM HEPES buffer and electroporated into cells using a Neon transfection system with 100 μL electroporation tips. The electroporation parameters for transfection of CD34+ cells included an electrical potential of 1600 V, pulse width of 10 msec over three pulses.

### Confocal Microscopy

G-CSF mobilized CD34^+^ HSPCs from three independent donors were thawed and plated in SFEM II media supplemented with 50 ng/mL TPO, Flt-3 ligand, and SCF. Cells were incubated overnight as described above before treatment with either 1^st^ generation CRISPR-AuNP or 2^nd^ generation CRISPR-AuNP, both synthesized as previously described with a fluorescently labeled ATTO550 trRNA. 6 hours after nanoparticle addition, cells were washed in DPBS twice before staining with NucBlue Live Cell Stain (Hoechst 33342, ThermoFisher) and CellMask Deep Red membrane stain (ThermoFisher) according to manufacturer’s protocol. Cells were washed twice more in DPBS and resuspended in DPBS with 2% FBS for acquisition on a Zeiss LSM 780 confocal microscope (Oberkochen, Germany). Images were acquired using a 63x oil immersion objective at 0.8x zoom for widefield capture. For fluorescence quantification, all cells in widefield images were manually counted. Cells with intact stained nuclei were included in the total live cell count. Cells with punctate ATTO550 fluorescence in the cytoplasm were designated as trRNA^+^. For each sample set, frequency was recorded as number of trRNA^+^ cells divided by the total in frame live cell count.

### Statistical analysis

Graphpad Prism (10.0.4; Boston, Ma, USA) was used to present all results as the mean ± SEM. Two-group comparisons used Student’s t-test. Multiple group comparisons used Tukey posthoc test and one-way ANOVA. The threshold for statistical significance was *α* < 0.05.

## Results/Discussion

### CRISPR Nuclease Loading on 1^st^ Generation CRISPR-AuNP

Cas9 is known to provide efficient indel formation in vivo upon DNA cutting relative to Cas12a.^44^ Comparatively, the CRISPR-AuNP developed by Shahbazi et. al.^45^, herein referred to as the 1^st^ generation CRISPR-AuNP, resulted in Cas12a and Cas9 CRISPR-AuNP with similar activity in vitro. Lower than expected Cas9 activity was originally attributed to lower efficacy of the specific gRNA designed to target the *CCR5* locus. However, further investigation with electroporated Cas9 RNP showed considerably higher functional indel formation relative to Cas12a at the *B2M* locus in Jurkat cells (Supplement 1,2C). Resulting Cas9 indel levels were also associated with higher B2M cell surface protein knockdown across all tested guides by flow cytometry, compared to Cas12a (Supplement 1), implicating another factor in the decreased Cas9 activity following CRISPR-AuNP formulation.

Direct measurement of Cas9 protein levels by SDS-PAGE densitometry after CRISPR-AuNP purification revealed undetectable levels of Cas9 nuclease compared to 3.4 Cas12a proteins/Au core (Figure 1A). Further, Cas12a AuRNP demonstrated dsDNA cutting after purification while Cas9 AuRNP cutting of dsDNA remained undetectable (Figure 1B). This indicated the decrease must be caused by loss of RNP during the initial two phases of nanoparticle assembly from AuRNA to AuRNP and contradicted existing results in cells, which showed Cas9 as able to deliver effective editing by Sanger sequencing and TIDE analysis. Further examination revealed the original 1^st^ generation CRISPR-AuNP formulation method left active RNP present in solution, that are possibly able to form polyplexes with PEI and associate with the particle surface electrostatically. Cas12a, on the other hand, was covalently attached to the Au core in 1^st^ generation CRISPR-AuNP, shown through considerable DNA cutting at all stages of assembly, and detectable nuclease levels by SDS-PAGE with additional purification (Figure 1), making the RNP loss during CRISPR-AuNP assembly specific to Cas9.

**Figure 1:**
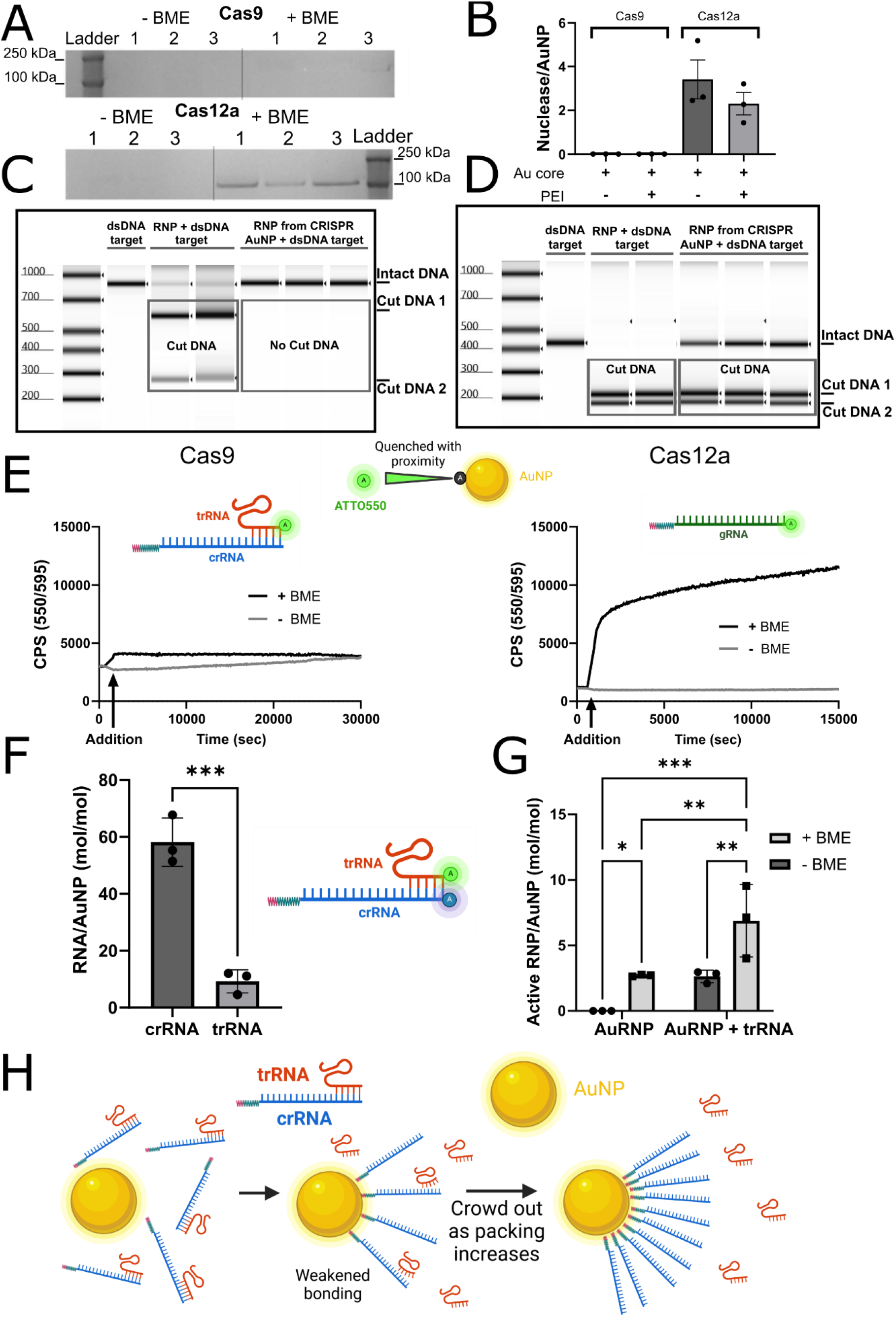
trRNA instability during 1^st^ generation CRISPR-AuNP assembly inactivates Cas9. A) Protein retention of 1^st^ generation CRISPR-AuNP after purification, staining with colloidal Coomassie, following each stage of synthesis. B) Quantification of SDS-PAGE by densitometry, converted to moles of nuclease per moles of Au core before and after PEI addition (18.3 nm Au core, n = 3). C) Cas9 and D) Cas12a DNA cutting assay for BME-released RNP from 1^st^ generation CRISPR-AuNP after purification, resulting in no detectable Cas9 activity. E) Quenched fluorescence recovery during kinetic release of fluorophore-labeled RNA from the Au surface. Notably, albumin containing buffer slows the release rate upon BME addition for Cas12a. F) Direct measurement of both crRNA-ATTO488 and trRNA-ATTO550 after purification and BME release, quantified against a standard curve. G) RNP activity is restored through re-addition of trRNA after particle purification and BME release, quantified by DNA cutting assay. H) Proposed schema of trRNA loss through de-duplexing during particle assembly. ATTO550 is depicted at the end of the trRNA, which is quenched in proximity to the gold core, indicating binding. After release with BME, a thiol competitor, the distance increases from the Au core, reducing quenching and recovering fluorescence. For figures, n = 3 technical replicates, bars represent means and error bars represent standard error of the mean. p = * < 0.033, ** < 0.0021, *** < 0.0002.

### Duplex RNA Loading on 1^st^ Generation CRISPR-AuNP

Cas9 and Cas12a both rely on thiolated gRNA anchors to bind the nuclease to the gold surface. As previously reported, the modifications required to adapt RNA to attach to a gold surface required a long chain spacer to buffer electrostatic repulsion between RNA molecules in addition to a thiol anchor. These modifications were not commercially available to form Cas9’s longer sgRNA, requiring the use of the crRNA/trRNA duplexing. Original testing of RNP by electroporation showed Cas9 maintained original activity when crRNA was modified on the 5’ end (unpublished data). One flaw is the duplex itself is held together by only 10 bp, with a melting point of 28 °C in our formulation conditions. Under low pH conditions required for AuRNA formulation, the melting point of duRNA is further reduced.^46^ Comparatively, Cas12a utilizes a shorter single guide (sgRNA, 41 bp) that could be thiolated on its 3’ end commercially. We used these two systems to approximate sgRNA vs the duplex method for these studies.

To investigate the binding of these RNA components to the Au core, ATTO550 was modified onto the trRNA for Cas9 and gRNA for Cas12a. These fluorophores are quenched by plasmon resonant energy transfer (PRET) within a 7 nm radius of the Au core.^47,48^ Kinetic analysis of Cas12a AuRNA showed effective PRET quenching after citrate binding, followed by fluorescence recovery after BME driven displacement of the gRNA’s thiol anchor (Figure 1C). Reaching a final concentration of 28 nM (60.7 gRNA/Au) and with a t_1/2_ of 3.02 hrs, it was clear that Cas12a gRNA was bound in a thiol-dependent monolayer to the gold surface. Furthermore, in the presence of 1% albumin and BME release the t_1/2_ increased to 12.78 hr, a reasonable estimated time for systemic in vivo delivery and uptake (Supplement 3). However, Cas9 duplex RNA (Figure 1C) labeled with trRNA/ATTO550 had a higher initial fluorescence signal, which suggests either weaker quenching or lack of binding to the Au core surface. After BME addition, a rapid and minor signal increase to an equivalent of 4.7 trRNA/Au was present after baselining, strongly suggesting poor thiol binding dependence. Additionally, a steady trRNA signal increase in the BME-free condition further indicated low stability of attachment to the surface.

With trRNA attachment stability a primary issue, crRNA-SH binding and duplex stability were called into question. The crRNA was labeled with ATTO488 for detection simultaneously with ATTO550 trRNA to confirm binding of both components. The purification of 1^st^ generation AuRNA formed with co-labeled duRNA revealed 9.2 ± 4.1 trRNA/Au core with 58.1 ± 8.5 crRNA/Au core (Figure 1D), leaving an unbalanced, trRNA-depleted surface. This loss is likely driven by low pH destabilization of the duRNA.^49^ As the fluorophores could disrupt the duplex, a DNA cutting assay was run with a bolus of trRNA added post-BME release to confirm trRNA loss as the primary cause of low Cas9 CRISPR-AuNP activity. This addback of tRNA increased Cas9 dsDNA cutting to a level akin to Cas12a CRISPR-AuNP (Figure 1E), at an equivalent of 4.3 ± 2.8 active RNP/Au core or 0.69 pmol/cm^2, when subtracting the non-supplemented trRNA activity. The non-trRNA and BME release activity represented 2.7 ± 0.3 RNP/Au core without trRNA addition, alone. The matching trRNA addition to non-trRNA + BME samples suggests some residual components were still present within the samples even after additional purification, but were sequestered into separate regions (on particle and free in solution) and therefore unable to become active until released and/or supplemented.

Further, Wei T. et. al. has shown that RNP aggregates stick to microcentrifuge tube walls at pH 3.8, which we confirmed by comparing the supernatant of Au core free reactions ± citrate, before and after purification, where trRNA fluorescence can be recovered by returning the supernatant to neutral pH (Supplement 4A).^50^ Additionally, we showed that trRNA alone was retained on the formulation tubes under similar conditions (Supplement 4B), though at lower levels. The low pH conditions used to facilitate RNA approach to the gold surface during synthesis also drives tube binding of RNA, allowing it to survive through the initial RNA purification stages where pH is maintained below pH 4.5. This low pH state had been maintained through the 1^st^ generation CRISPR-AuNP formulation, with neutrality only occurring during PEI addition. This supported our hypothesis that in vitro editing observed with the 1^st^ generation CRISPR-AuNP was likely through trRNA release from the tube rather than fully functional Cas9 CRISPR-AuNP assembly. The additional purification required for this in depth testing involves a tube change before pH neutralization, excluding tube bound trRNA. This made the re-addition of trRNA a requirement for CRISPR cutting activity, being more apparent in subsequent results.

The possibility of Cas9 rebinding after trRNA supplementation was determined kinetically (Supplement 5) and by SDS-PAGE after trRNA + Cas9 Au-crRNA incubation for 1 hr. However, this resulted in no quenching of trRNA-ATTO550, nor enhancement of Cas9 retention in SDS-PAGE. The hypothesized reason for this low binding is derived from best case surface saturation and volume fill calculations (Supplement Table 2), which show lower surface densities are required for the duplex gRNA vs crRNA-SH alone. For Cas9, the absolute surface loading limit is near 69 RNP for an 18 nm Au core using best case volume fill calculations, and 17 RNP/Au core for surface projection ellipsoid packing. After crRNA saturation, the packed surface excludes RNP rebinding. With these conclusions in mind, a method to achieve ideal packing for the RNP alone is critical to making a maximally loaded system.

### Stabilizing duRNA by Preformation of CRISPR RNP

Due to the low affinity of cr/trRNA duplex during the 3.0 pH AuRNA coating, as well as crRNA packing disfavoring RNP binding post AuRNA formation, methods for increasing Cas9 gRNA duplex affinity and surface chemistry were investigated. Research by O’Reilly et al. showed increased Cas9 nuclease melting temperature after addition of duplexed RNA, indicating enhanced RNP stability.^51^ We hypothesized that this increased stability is reflected in both the protein and duplex RNA. Therefore, using preformed RNP would increase the binding stability of the duplex as well as restricting the surface to only binding full RNP, instead of forming RNA-only layers. However, even when including a neutral PEG linker, RNA and citrate coated Au cores require low pH to neutralize their negative charge and achieve binding. A pre-formed RNP also carries negative charge (Cas9 = -5-10 mV), requiring less harsh coating pH relative to RNA but also risks permanent loss of activity.^52^ Therefore, we investigated the retention and activity of both Cas9 and Cas12a RNP from neutral pH to pH 2.5, where denaturation is assured, using a DNA cutting assay (Figure 2B). After AuRNP coating and purification, RNP activity showed an IC50 citrate buffer pH of 3.23 for Cas9, while Cas12a was more stable at pH 3.01. The peak activity for Cas9, corresponding to a combination of AuRNP binding and pH resistance, was pH 3.8. To unify particle formulation, pH 3.8 was also employed for Cas12a, whose maximum activity peak was observed between pH 3-3.2.

**Figure 2:**
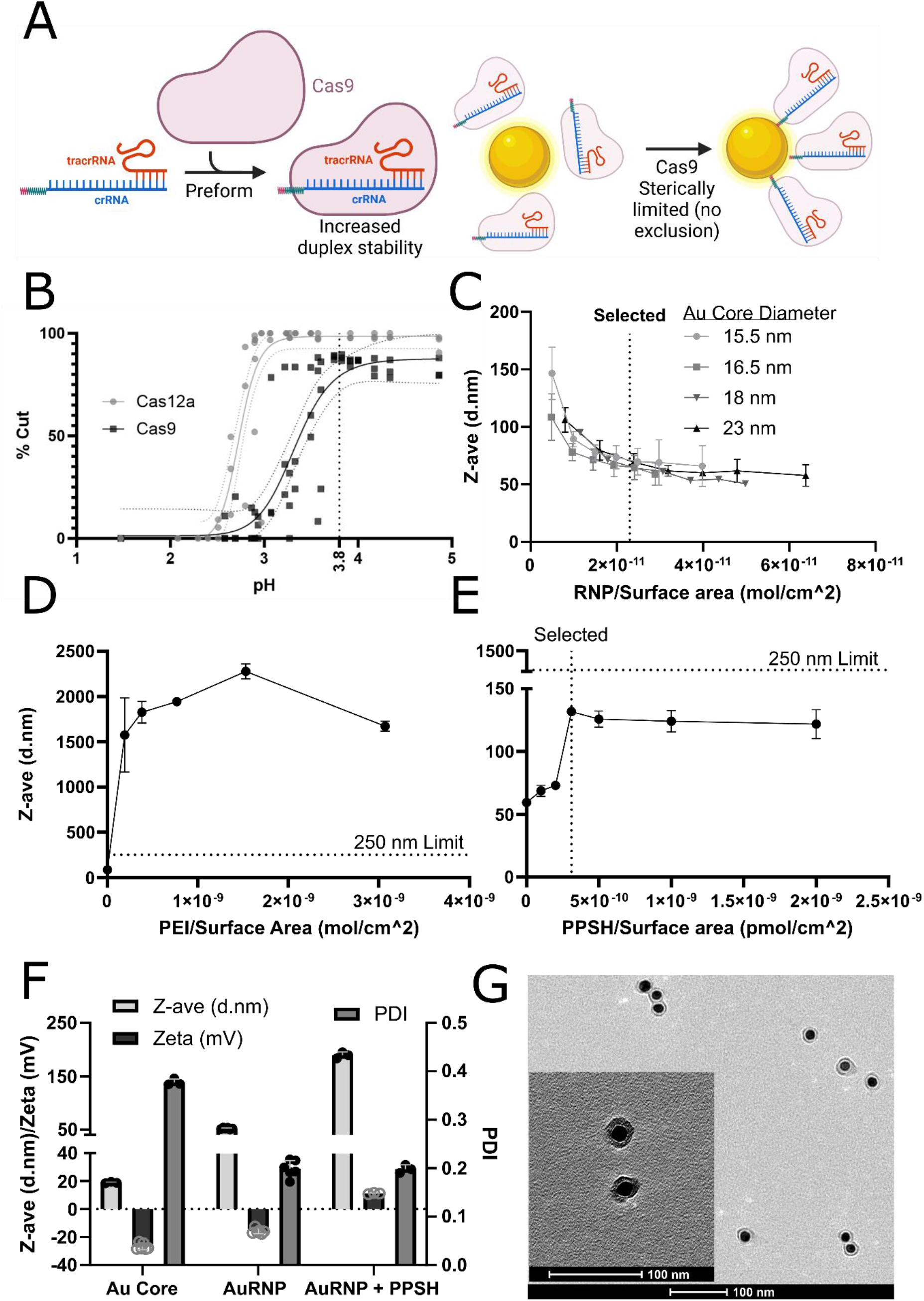
Preformed RNP is preserved through low pH formulation, increasing loading and activity of 2^nd^ generation CRISPR-AuNP. A) Schematic for the preformation and coating of AuRNP, utilizing Cas9’s increased affinity for cr/trRNA to maintain duplex and extant volume to limit overpacking. B) DNA cutting assay of 2^nd^ generation Cas9 AuRNP formed with a range of citrate pH stocks and purified by centrifugation, then released from the particle surface with BME. C) DLS measurements of AuRNP size vs preformed RNP addition ratio for a range of gold core sizes. D) DLS showing destabilization of AuRNP coated with PEI. E) DLS of differing PPSH coating levels that stabilizes near 120 nm. F) Particle characteristics of 2^nd^ generation CRISPR-AuNP at each stage of synthesis, including the zeta potential. G) 0.1% uranyl acetate-stained 2^nd^ generation CRISPR-AuNP with complete coating. For figures, n = 3 technical replicates. Bars represent means, error bars represent standard error of the mean.

To maximize Au surface loading and formulation efficiency, the RNP concentration vs Au core surface area was investigated by formulating differing Au core diameters against several RNP doses. The Au core surface area was calculated assuming spherical cores with the diameter determined by TEM (n = >10 particles measured per batch). AuRNP generated at low coating conditions resulted in aggregation (Figure 2C), with a minimum stable loading dose equivalent to 3.5 pmol RNP/cm^2. Additionally, increased coating was shown to reduce the particle size towards 50 nm, approaching the idealized monolayer radius (d_H_ = 20 nm Au core + d_H_ 2*12 nm Cas9 RNP + 2*2 nm PEG spacer = 48 nm). Additionally, as RNP density increased, zeta potential neutralized, from a -30 mV potential, similar to AuRNA and bare gold, to between -20 to -15 mV (Figure 2D), a charge decrease representative of the positive Cas9 nuclease neutralizing 33-50% of the RNA charge, with expected neutralization near 30%. The 23 pmol/cm^2 formulation dose of RNP was selected for the <10% trend in Z-ave and zeta potential with a minimum in polydispersity index (PDI) (Supplement 6).

### AuRNP Coating with PEI Derivatives

Our 1^st^ generation CRISPR-AuNP included 2K, branched chain PEI to create positive charge, making CRISPR-AuNP electrostatically attractive to negatively charged plasma membranes and promote endosomal escape. However, addition of this PEI to AuRNP assembled with preformed RNP resulted in particle aggregation to >1 µm diameter (Figure 2D). It is hypothesized that reduced zeta potential of the 2^nd^ generation AuRNP lowered electrostatic attraction to PEI, resulting in unfavorable layer formation kinetics and interparticle adhesion as colloidal stability decreased. To overcome this, PEI was grafted with thiolated 2k Mn PEG (PPSH), with the PEG-SH acting as an anchor to the gold surface. We hypothesized that thiol-anchoring increases colloidal stability through steric repulsion and could increase biocompatibility for future in vivo testing. Coating at 32 ug PPSH/40 ug Au core (Figure 2E), resulted in an 87.7 ± 5.9 nm CRISPR-AuNP (0.198 ± 0.012 PDI and 3.6 ± 0.9 mV, n = 5) with a +5 mV charge (supplement 9). Further, these particles were stable against media and serum conditions (Supplement 7) suggesting biocompatibility for in vivo administration. This structure was designated as 2^nd^ generation CRISPR-AuNP.

### Quantification of RNP on 2^nd^ Generation CRISPR-AuNP

Fully formulated 2^nd^ generation CRISPR-AuNP showed 39.6 ± 7.0 Cas9 RNP/AuNP by SDS-PAGE when released with BME (Figure 3A), and dsDNA cutting assay demonstrated >50 active Cas9 RNP/AuNP using the standard curve method (Figure 3B). This formulation of 2^nd^ generation CRISPR-AuNP also resulted in significantly improved Cas12a loading and retention by SDS-PAGE (Figure 3C) and dsDNA cutting activity (Figure 3D) as compared to 1^st^ generation CRISPR-AuNP (Figure 1A and 1B). The lower Cas12a loading level may be driven by the higher pH selected for formulation, which was based on Cas9 stability (Figure 2B). Designing the loading pH specifically to Cas12a’s increased pH stability may increase the loading level.

**Figure 3:**
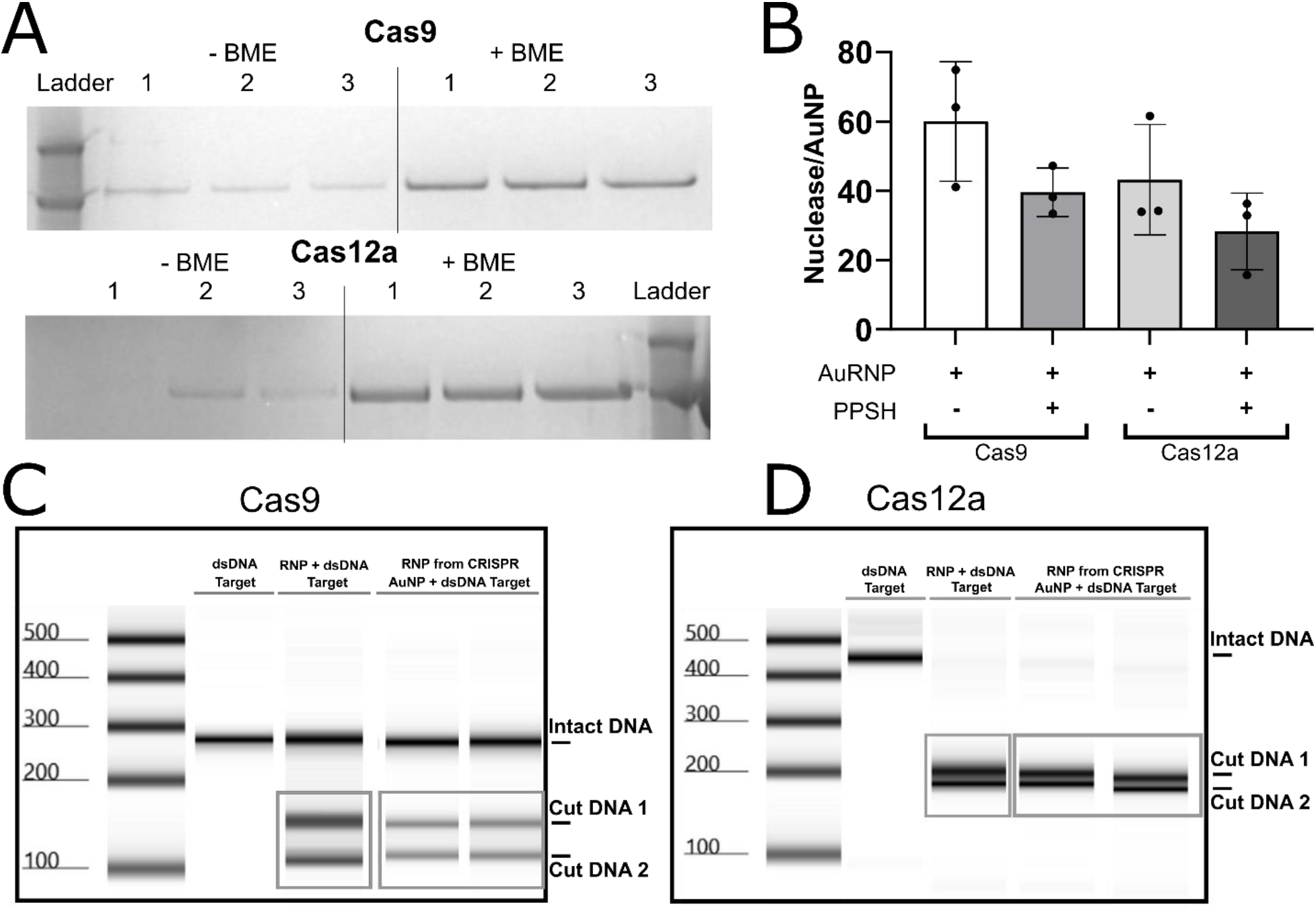
2^nd^ generation CRISPR-AuNP improves active RNP loading for Cas9 and Cas12a. A) SDS-PAGE of purified 2^nd^ generation CRISPR-AuRNP before and after BME addition. B) Quantification of SDS-PAGE by densitometry, converted to mol nuclease per mol Au core before and after PPSH addition (19 nm Au core, n = 3 technical replicates). C) Cas9 and D) Cas12a DNA cutting assay after BME release of 2^nd^ generation AuRNP. Bars represent mean, error bars represent standard error of the mean.

### B2M CRISPR Knockdown in HSPCs

With stable and active RNP loaded 2^nd^ generation CRISPR-AuNP, direct comparison to the 1^st^ generation CRISPR-AuNP was developed. To allow genomic and translational quantification of gene editing, the human *B2M* locus was selected for gene editing. B2M is a surface expressed protein in nucleated cells. The *B2M* exons were screened for all possible Cas9 gRNAs, then the >100 possible guides were winnowed using Chen W et. al.’s method for determining a target site’s ability to perform homology directed repair (HDR), with the top 20 selected for screening.^53^ Electroporation of RNP into Jurkat cells was evaluated by flow cytometry for loss of cell surface B2M protein and DNA sequencing to determine editing rates in genomic DNA (gDNA) (Supplement 1,2). Of the guides tested, Cas9 guide 4, targeting Exon 2 in a region vital to B2M interaction with major histocompatibility class (MHC) I protein, knocked down B2M expression 54.7 ± 20.6% and was associated with 65.0 ± 24.0% gene editing (defined by altered DNA sequence reads relative to the cut site vs total reads). The high variation stemmed from one donor having consistently low gene editing, with the trend in guide activity holding for all three donors evaluated. Guides which significantly suppressed B2M surface expression targeted either the promoter region or exons 1/2, while even high gene editing at the gDNA level in exons 3/4 were associated with little to no knockdown of B2M protein expressed on the cell surface. Guide 4 was selected for its high B2M protein knockdown which permitted rapid determination of Cas9 editing efficiency in living cells by flow cytometry to facilitate further CRISPR-AuNP development.

Having selected guide 4, the RNP dose required to achieve effective editing, assuming cytoplasmic delivery of RNP, was quantified. To this end, a range of RNP was electroporated into HSPCs and compared against mock treatment. A dose of 100 pmol RNP demonstrated the highest stable gene editing at 43.9 ± 17.9% B2M^low/-^ by flow cytometry and 47.8 ± 15.3 % edited reads by DNA sequencing (Supplement 8), and 20 pmol demonstrated the lowest detectable gene editing with statistical power at n = 3 biological replicates by DNA sequencing. These doses were selected for initial CRISPR-AuNP study doses.

### HSPC Delivery of Cas9 by CRISPR-AuNP

Targeting a 100 pmol dose of RNP, the corresponding CRISPR-AuNP dose was calculated based on SDS-PAGE bound RNP/Au core. HSPCs showed 73.7 ± 6.1% viability (Figure 4A) after 3 days incubation with 100 pmol RNP equivalent dose of 2^nd^ gen CRISPR-AuNP (45.6 μg of 15.5 nm Au core). This treatment produced a non-statistically significant trend in gene knockdown that was below the expected value from increased RNP loading density (Figure 4B,C). Higher doses were not pursued as this risked sub 70% viability, a level known to limit both in vitro efficiency targets and in vivo compatibility.

**Figure 4:**
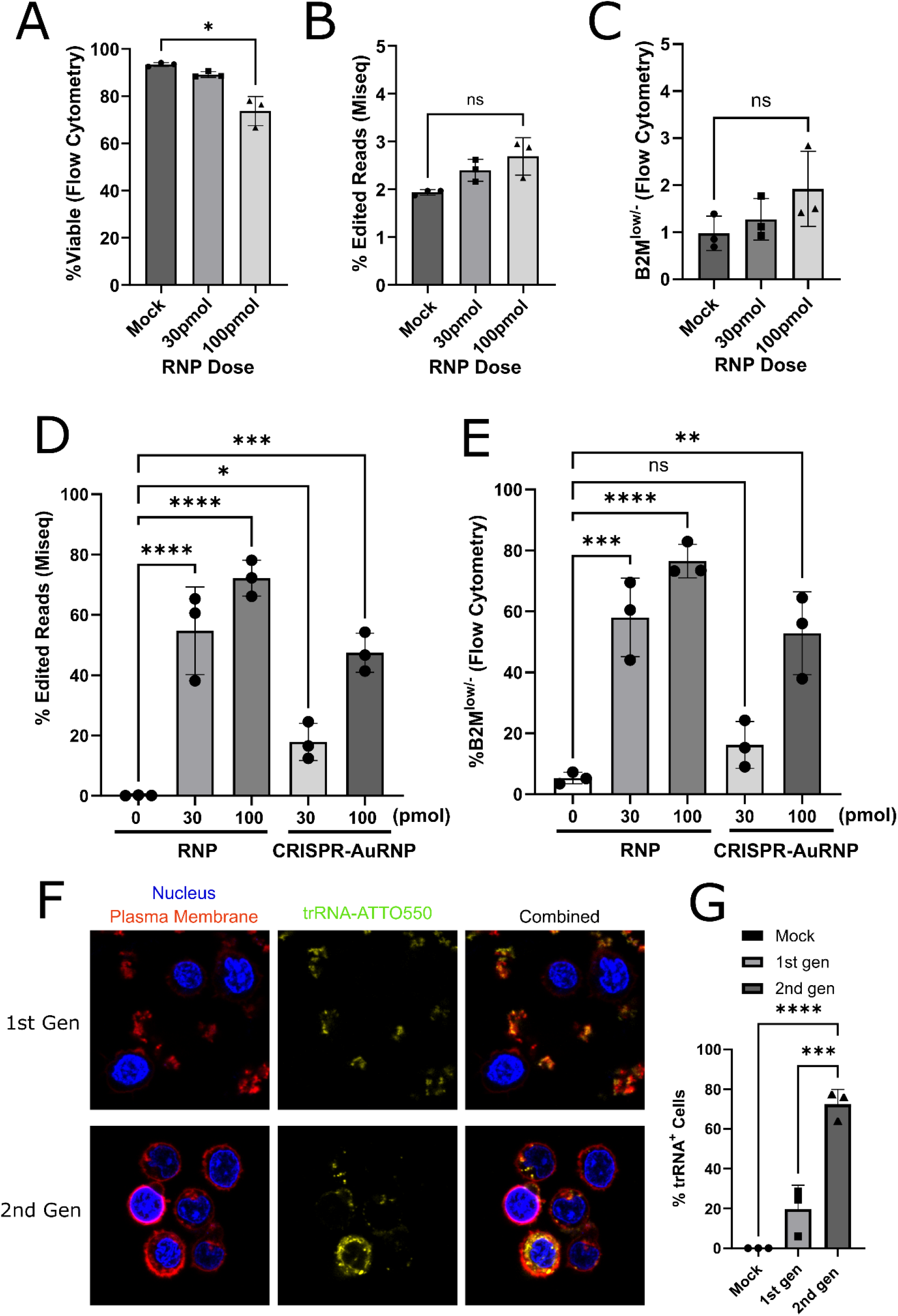
Cellular treatment with 2^nd^ generation CRISPR-AuNP determines optimal dose and indicates endosomal entrapment. A) Live/dead flow cytometry viability measure of primary human CD34^+^ HSPCs after treatment with 2^nd^ generation CRISPR-AuNP at two RNP doses. B) Gene editing rates determined by DNA sequencing, quantifying the % of reads which contained a DNA alteration within 40 bp of the cut site, resulting from 2^nd^ generation treatment of CD34^+^ cells. C) B2M protein knockdown rates determined by flow cytometry of CD34^+^ cells after 2^nd^ generation CRISPR-AuNP treatment. D) Gene editing rates from BME released RNP, electroporated into CD34^+^ cells, from 2^nd^ Generation AuRNP quantified as % DNA reads with edits within 40 bp of the cut site. E) B2M protein knockdown rates determined by flow cytometry of CD34^+^ cells after electroporation of 2^nd^ generation BME released RNP. F) Confocal microscopy images of 1^st^ and 2^nd^ generation CRISPR-AuNP treated CD34+ cells. Nuclear staining (blue) and red membrane stain were used to delineate cytosol; CRISPR-AuNP were tracked with trRNA-ATTO550. F) Quantification of percentage of CD34+ cells after confocal imaging that were positive for cytoplasmic trRNA-ATTO550 staining. For figures n = 3 independent donors, bars represent mean, error bars represent standard error of the mean, p = *<0.033, **<0.002, ***<0.0002, ****<0.0001.

To test RNP loaded onto CRISPR-AuNP for in vitro activity, 2^nd^ generation CRISPR-AuNP were formulated to AuRNP stage and bound RNP were electroporated into primary HSPCs (Figure 4D,E). Electroporation of BME-displaced RNP from CRISPR-AuNP at a targeted 100 pmol dose was compared against free RNP. The calculated 100 pmol dose resulted in equivalent gene editing to 30 pmol free RNP, 1/3 the expected RNP loading level by SDS-PAGE calculations. As controls exposed to BME showed no alteration in gene editing rates, this suggests some decrease in activity on the AuNP, setting the active RNP loading at ∼16 RNP/Au core. While this dose is lower than expected, the 2^nd^ Generation CRISPR-AuNP contains a sufficient active RNP dose to produce statistically significant genetic editing using the B2M model.

With an active RNP dose sufficient to achieve genetic editing, assuming cytoplasmic delivery, we further explored the CRISPR-AuNP uptake mechanism in HSPC. 2^nd^ generation CRISPR-AuNP were tracked via confocal microscopy of ATTO550-trRNA labeled CRISPR-AuNP (Figure 4E). After 6 hr of treatment, both 1^st^ and 2^nd^ generation CRISPR-AuNP demonstrated punctate staining within the cytoplasm, supporting endosomal internalization. When visually quantified, a high overall accumulation was apparent in HSPC treated with 2^nd^ generation CRISPR-AuNP, whereas 1^st^ generation CRISPR-AuNP-associated fluorescence was associated with extracellular PEI accumulation (Figure 4G). However, there was no evidence for endosomal escape of trRNA associated with either 1^st^ or 2^nd^ generation CRISPR-AuNP, indicating endosomal entrapment.

As 1^st^ generation CRISPR-AuNP does result in gene editing, its lack of clear endosomal disruption here is possibly driven by the visible sequestration of trRNA in aggregates of PEI, seen floating between cells, suggesting instability under imaging conditions. While the 2k PEI within these aggregates has endosomal escape potential, it cannot enter cells in the state seen here and the lack of uptake and release being either a byproduct of the imaging conditions or any internalization of smaller fragments being washed out by the bright aggregates. The clear accumulation of trRNA in the aggregates suggests the existing editing rate of 1^st^ generation CRISPR-AuNP may still have been maintained by similar electrostatic association between trRNA to the PEI, even after initial loss from the duRNA during AuRNA formulation.

2^nd^ generation CRISPR-AuNP are seen to internalize with trRNA, but are unable to escape the endosome. Possibly driving this decreased endosomal disruption is the high level of PEG to PEI in the PPSH, supported analysis by Mariani et al.^54^ PEGylation of PEI is shown to limit its endosomal escape through disruption of direct membrane association, a process required to form pores in swelling endosomes. As the particles here use 2k MW PEI grafted at 10% of amines with 2k PEG, the ratio is 1:5 PEI:PEG by weight. This sterically limits PEI’s interaction to semi-ridged endosomal membranes, lowering pore formation. Of note, our results support Mariani’s conclusion that the simplistic proton sponge hypothesis is insufficient to explain PEI activity. PEGylation does not alter proton affinity nor significantly reduce proton capacity, being easily adjusted for by dose alone. Reduction of the PEG coating is currently under investigation, along with incorporating other endosomal disrupting agents.

## Conclusions

We have shown Cas9’s duRNA alone is unable to maintain stability through AuRNA formulation of the 1^st^ generation CRISPR-AuNP, substantially reducing the activity after two washes. This loss results in no effective DNA cutting activity in tubo after purification, where the 3^rd^ centrifugation wash and tube transfer removes surviving trRNA. The remaining activity of Cas9 observed in the original publication was likely driven by non-specific retention and PEI binding of the trRNA, but this approach is incompatible with in vivo stability.

Utilizing Cas9’s biophysical interactions between nuclease and duRNA to increase stability, preforming the RNP resulted in 2^nd^ generation CRISPR-AuNP with Cas9 present at an average of 16 active RNP/AuNP by electroporation and 39.6 ± 7.0 RNP/AuNP by SDS-PAGE. This approach also increased loading of Cas12a. However, loading of preformed RNP required altered PPSH coating to achieve particle stability, which was associated with increased internalization but also considerable endosomal entrapment in HSPCs. We hypothesize this is directly caused by heavy PEGylation of the PEI endosomal escape agent, limiting the ability of PEI to release and associate with the endosomal membrane. We propose further investigation of alternate polymer structures to overcome this block, as well as utilizing alternative endosomal disruption agents to confirm endosomal escape as the primary block to CRISPR-AuNP activity.

In conclusion, direct understanding of the interaction of CRISPR systems to the AuNP-anchored portion of the RNP determines its stability when transferred in vitro. Further, leveraging the interaction between the protein and RNA can be used to increase duRNA stability and control loading level on the surface of AuNP. This approach shows increased loading, stability, and uptake of the resulting CRISPR-AuNP system, while maintaining the redox sensitive release mechanism essential to delivery. This improved compatibility is essential to utilization of novel Cas9 and other nuclease systems for future delivery and in vivo gene therapy.

## Acknowledgments

We thank healthy donors who submitted to mobilization and leukapheresis collection. We thank the following departments and labs within the Fred Hutchinson Cancer Center; the Cooperative Center for Excellence in Hematology who performed blood product extraction, donor organization and cellular purification. The Electron microscopy core for access and instruction for TEM analysis of CRISPR-AuNP. The Cellular Imaging core for confocal microscopy assistance. The Flow Cytometry core for access, maintenance, and training on the FACSCelesta flow cytometer. This work was primarily supported by the following grants: R01 AI167009 and R01 AI158728, and funds through J.E.A. from the Fred Hutch including Development and Evergreen awards, the Bill and Melinda Gates Foundation, and the Kuni Foundation. This research was also supported by the NIDDK Cooperative Center of Excellence in Hematology Grant U54 DK106829. All shared resources used in this study were supported by the NIH/NCI Cancer Center Support Grant P30 CA015704. J.E.A. is The Fleischauer Family Endowed Chair in Gene Therapy Translation and a co-inventor on U.S. Patent WO2018226762A1 entitled “Genomic Safe Harbors for Genetic Therapies in Human Stem Cells and Engineered Nanoparticles to Provide Targeted Genetic Therapies.”

## Author Contributions

D.D.L. drafted the manuscript as well as designed, planned, executed, interpreted, and analyzed experiments regarding 1^st^/2^nd^ generation CRISPR-AuNP, physiochemical analyses of Cas9 and Cas12a, DNA cutting assay, loading optimization, TEM and CRISPR-AuNP characteristics. K.S.V.G. conceived the 2nd generation CRISPR-AuNP design, designed, planned, executed, interpreted, and analyzed SDS-PAGE, electroporation, Jurkat and primary HSPC experiments in vitro, as well as *B2M* guide discovery and analysis. R.A.C. conceived the DNA cutting assay and designed, planned, executed, analyzed, and interpreted confocal microscopy, electroporation and primary HSPC experiments in vitro. M.E.C. and Y.J. performed DNA cutting assays, pH studies, flow cytometry, TEM, and in vitro Jurkat *B2M* guide selection and primary HSPC experiments and drafted methods/figures for the manuscript. J.M.P.C. performed Lindel analyses to identify ideal *B2M* guide RNA sequences and supported flow cytometry and gene editing analyses. J.E.A. funded the study and contributed to conception, design, analysis, and interpretation of data. All authors contributed to writing the manuscript in some capacity, as well as reviewed and edited the manuscript.

## Supplement

**Supplement 1:**
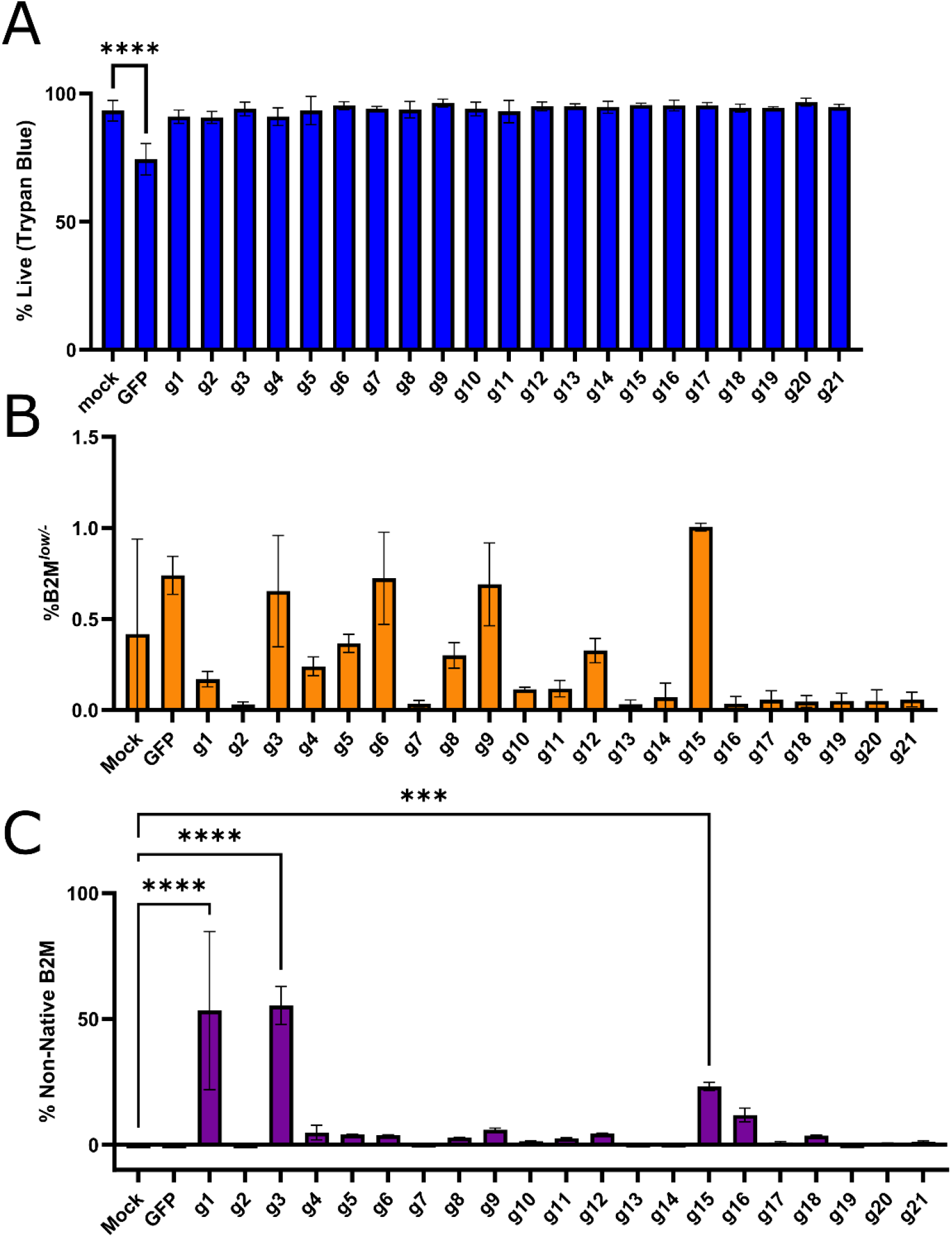
*B2M* Cas12a guide search. A) Viability of Jurkat cells electroporated with selected Cas12a guide targets as RNP using a Neon Electroporation system. B) Resulting % B2M^low/-^ Jurkat cells determined by flow cytometry for each guide tested. C) % edited DNA sequence reads (defined by indels or substitutions within 40 bp of cut site) for each guide. p = * < 0.033, ** < 0.0021, *** < 0.0002. n = 3 technical replicates, bars represent mean, error bars represent standard error of the mean

**Supplement 2:**
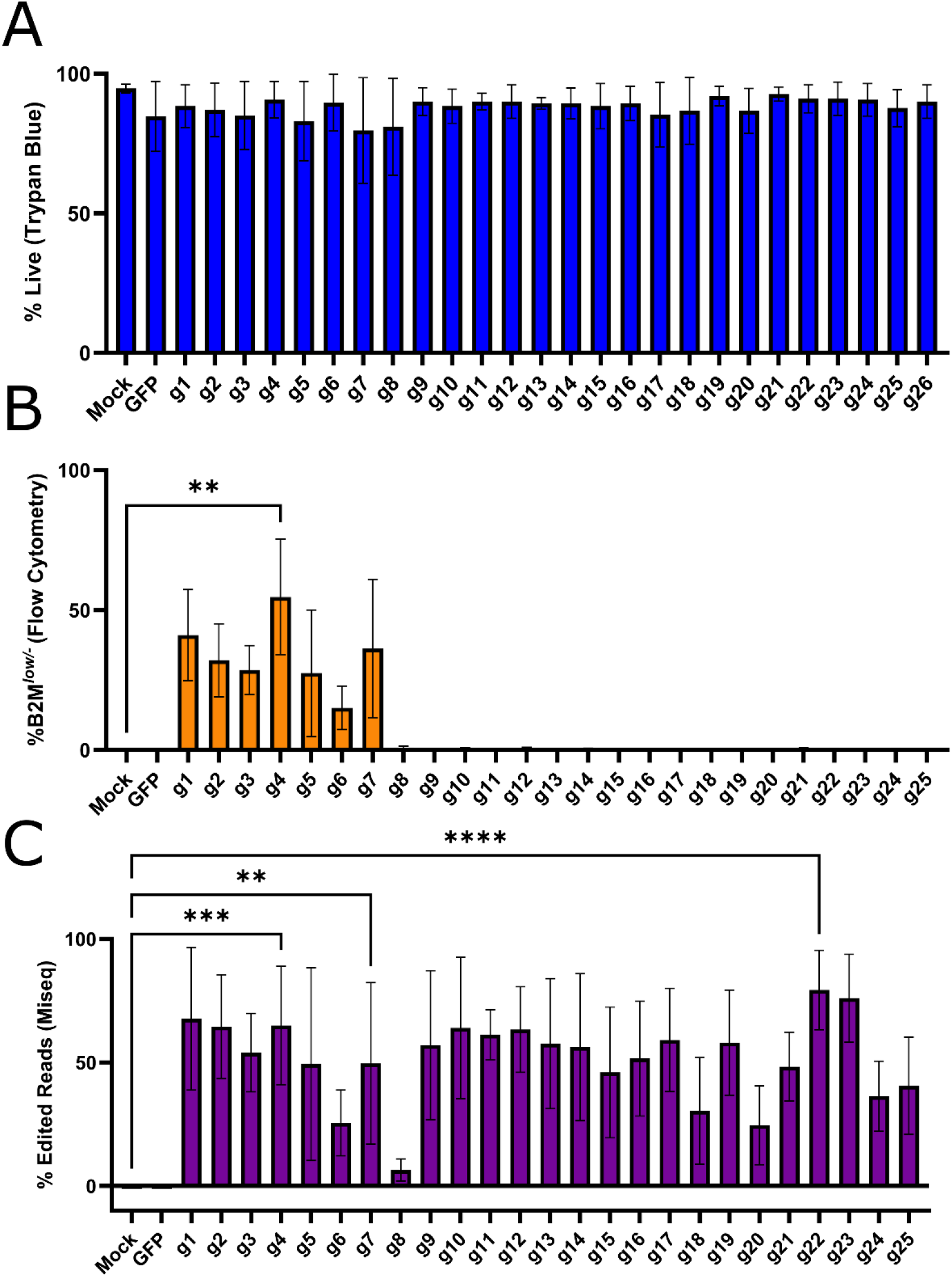
*B2M* Cas9 guide search. A) Viability of Jurkat cells electroporated with selected Cas9 guide targets as RNP using a Neon Electroporation system. B) Resulting % B2M^low/-^ Jurkat cells determined by flow cytometry for each guide. C) % edited DNA sequence reads (defined by indels or substitutions within 40 bp of cut site) for each guide. p = * < 0.033, ** < 0.0021, *** < 0.0002. n = 3 technical replicates, bars represent mean, error bars represent standard error of the mean.

**Supplement 3:**
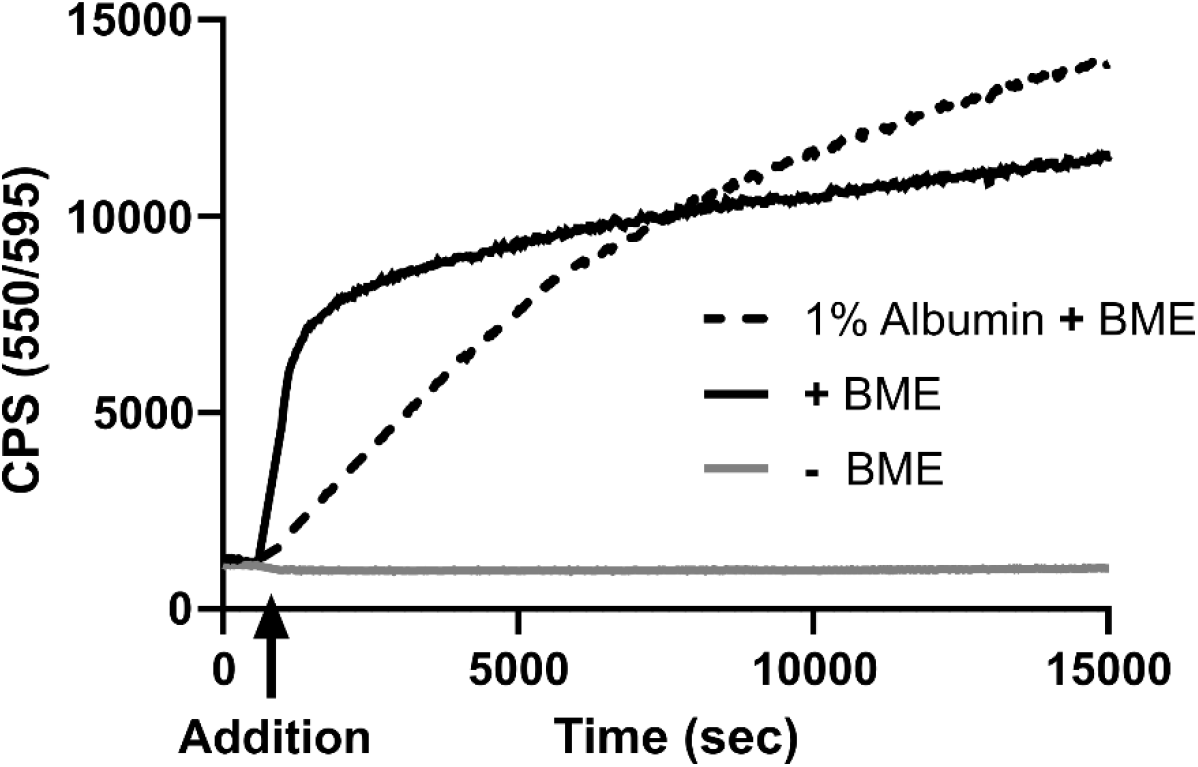
Kinetic release of 1^st^ generation Cas12a gRNA in the presence of albumin. Fluorescence recovery trace of gRNA-ATTO550 after BME release in the presence of albumin. Interactions of the albumin and particle slow the release rate to biologically relevant levels when BME is present at doses included in standard media.

**Supplement 4:**
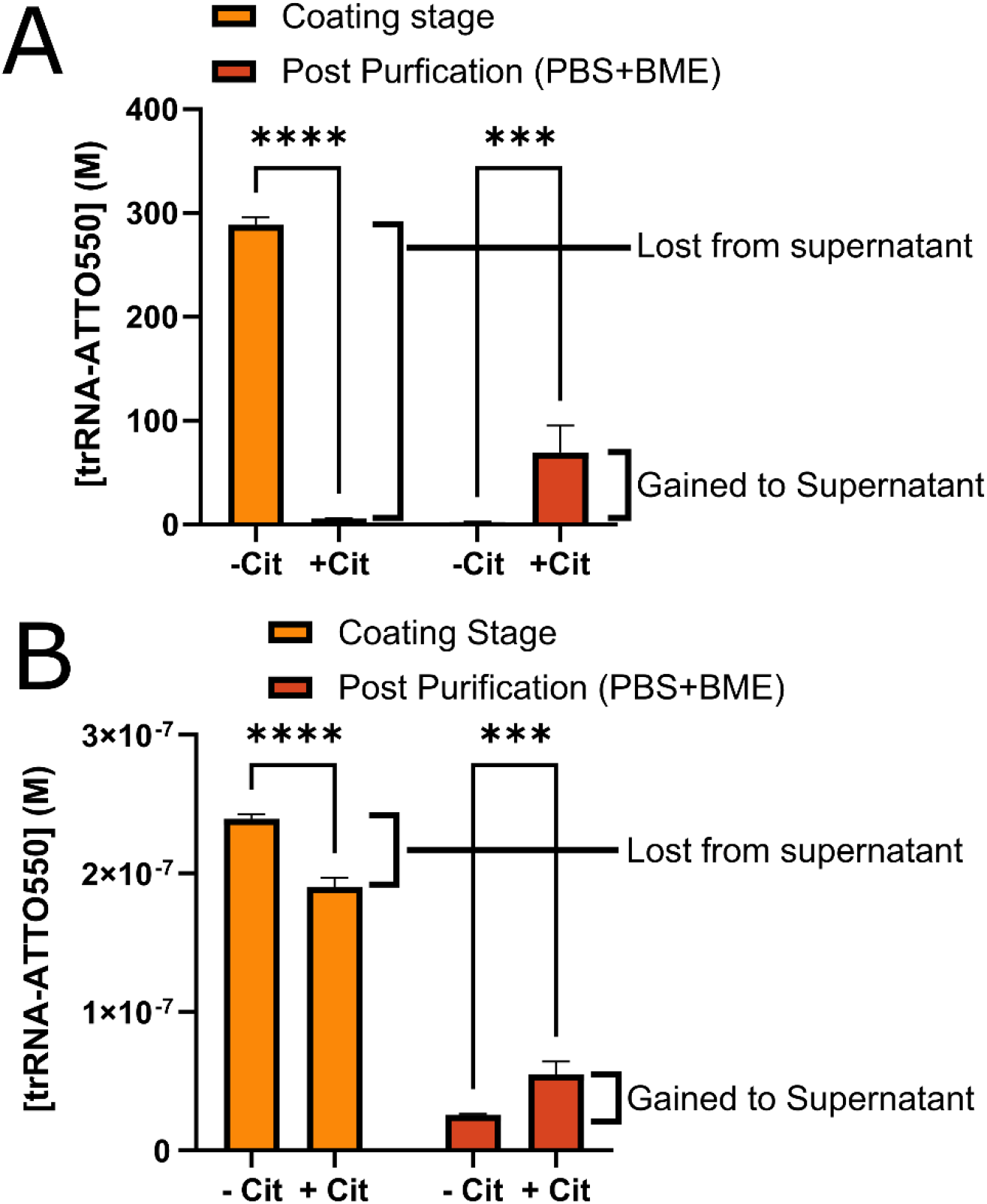
Retention of RNP and trRNA at low pH. A) trRNA-ATTO550 traced preformed RNP through 2^nd^ generation coating. Citrate negative samples had most RNP removed after the first spin, while no fluorescence was present in the citrate, and therefore low pH, sample. A portion of this can be recovered by adding BME and PBS to the citrate tube, which likely released RNP present in a pelleted aggregated state in the tube or on the wall. B) trRNA-ATTO550 traced duRNA through the standard 1^st^ generation coating. Some portion is lost for the supernatant at low pH and can be recovered through neutralizing and BME addition. For figures, n = 3 technical replicates, bars represent mean, error bars represent standard error of the mean, p = * < 0.033, ** < 0.0021, *** < 0.0002.

**Supplement 5:**
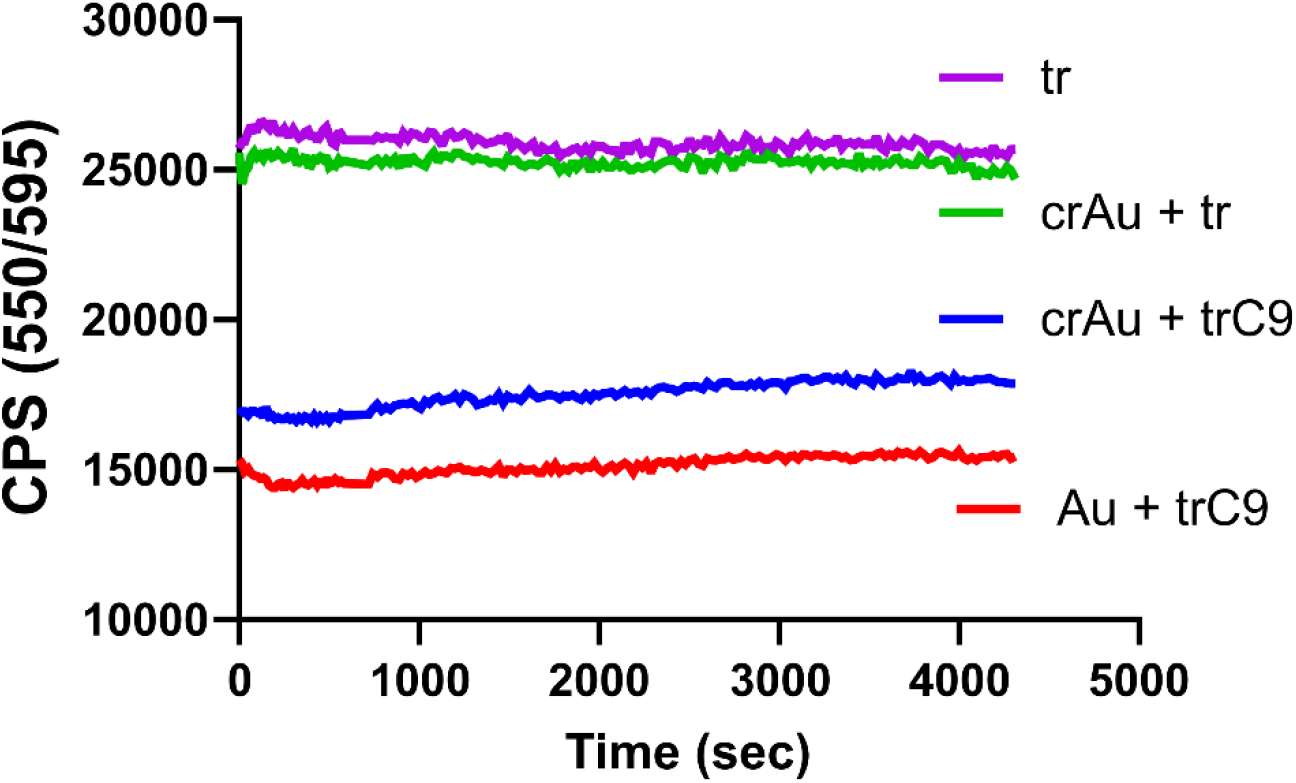
Kinetic interaction of Au-crRNA + tracRNA-Cas9. Fluorescence kinetic trace of trRNA-ATTO550 combined with Cas9 after addition to a crRNA-only modified Au core. Interactions which bring the trRNA within 7 nm of the Au core should result in PRET, however, no statistically relevant trend was present in any samples. This suggests a lack of interaction between the gold core and the fluorophore, with and without Cas9 as a mediator, after surface coating with crRNA at high density.

**Supplement 6:**
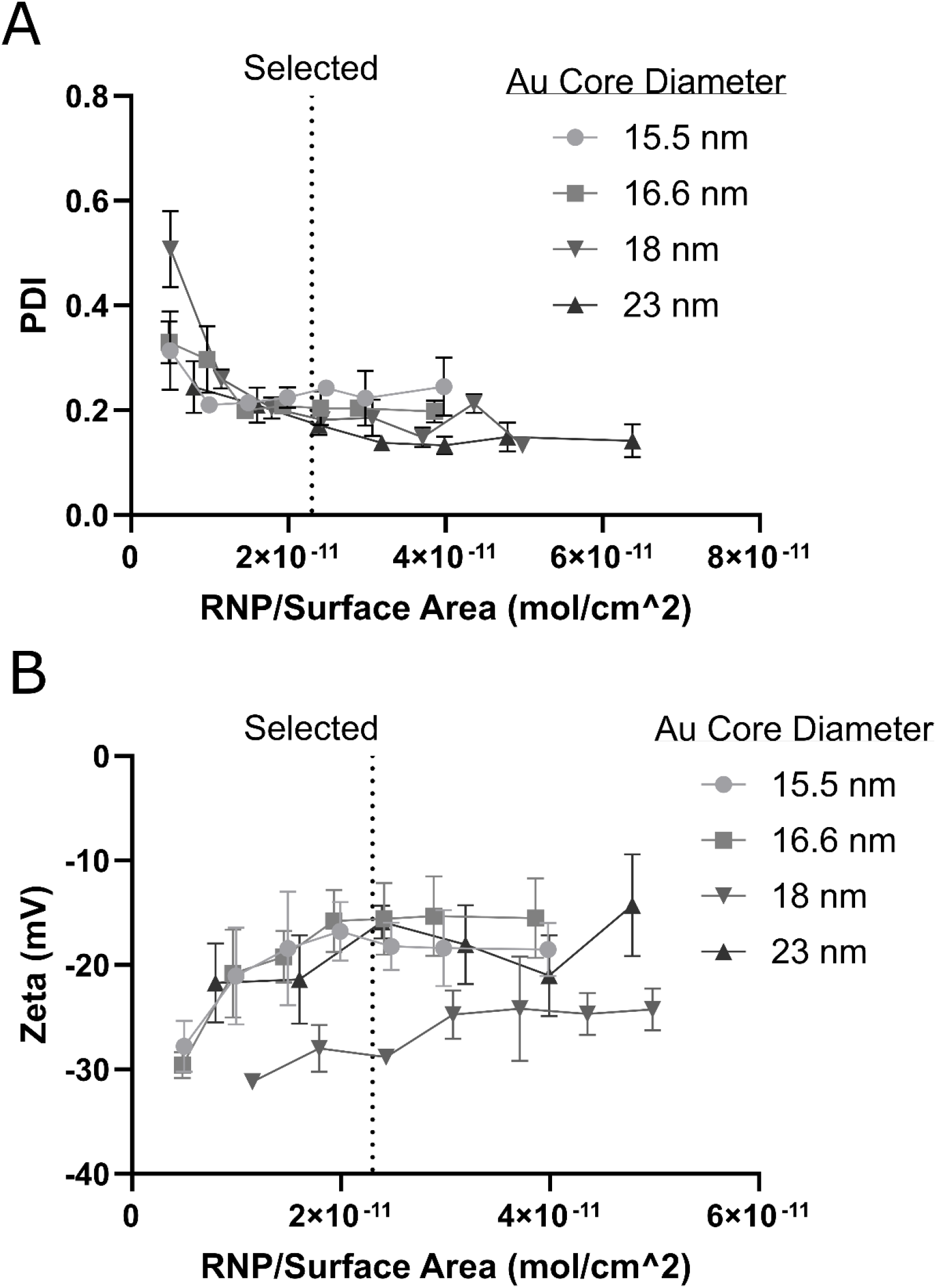
2^nd^ generation RNP coating dose, PDI/Zeta potential. A) PDI for several Au core sizes across a range of preformed RNP coating densities. B) Zeta potential for 2^nd^ generation AuRNP in 10 mM HEPES buffer at pH 7.4. For figures, n = 3 technical replicates, error bars represent standard error of the mean.

**Supplement 7:**
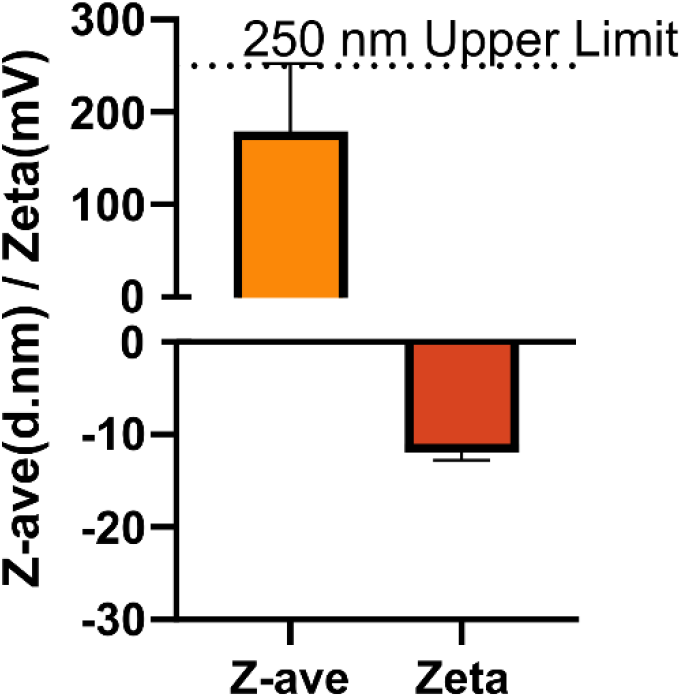
Serum stability of 2^nd^ generation CRISPR-AuNP. A) DLS reading of 2^nd^ generation CRISPR-AuNP in 10% serum. For figures, n = 3 technical replicates, error bars represent standard error of the mean.

**Supplement 8:**
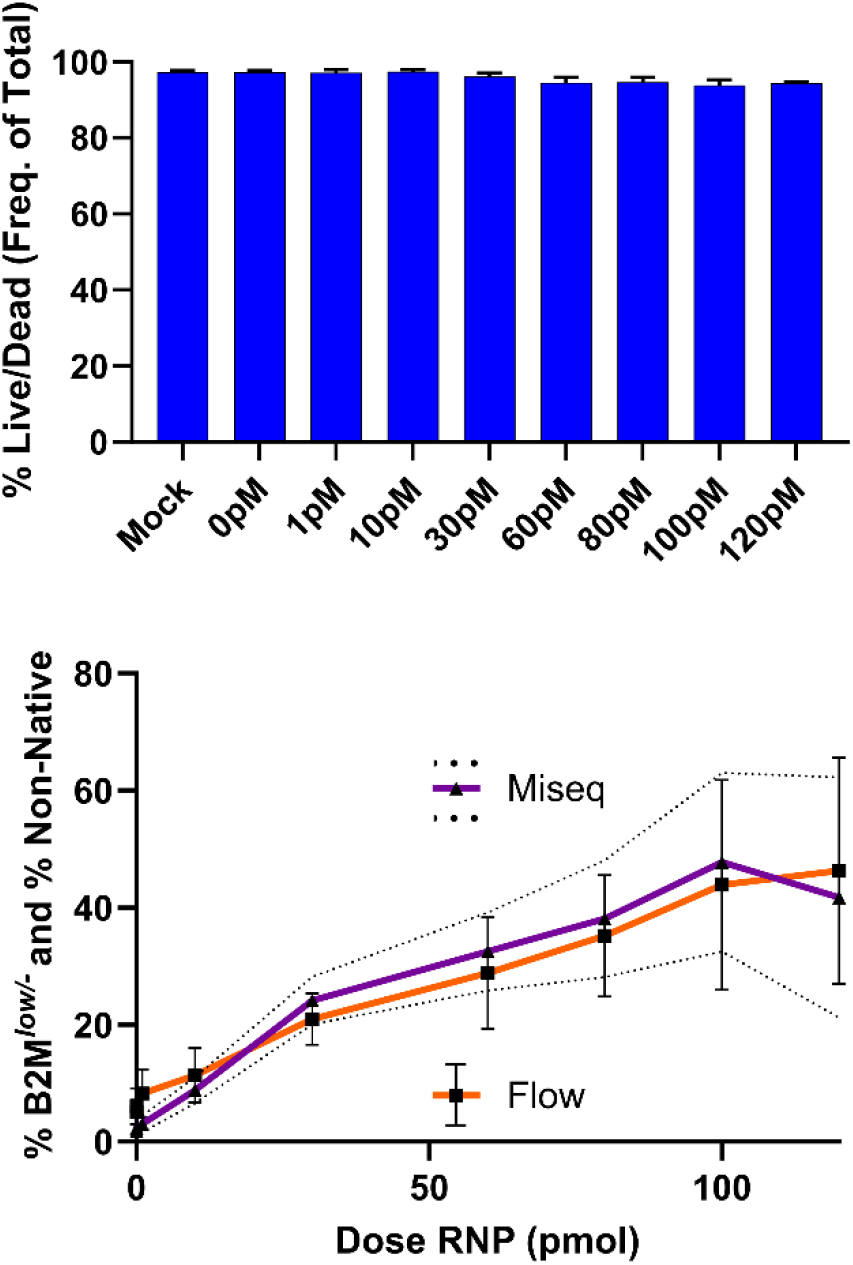
*B2M* Cas9 guide 4 electroporation dose curve. A) Viability of primary human CD34+ HSPCs electroporated at varying doses of RNP using a Neon Electroporation system. (n = 3 independent donors, bars represent mean, error bars represent standard error of the mean). B) Resulting % edited DNA sequence reads (defined by indels or substitutions within 40 bp of cut site) and % B2M^low/-^ HSPCs determined by flow cytometry. Dotted lines are standard error of the mean for gene editing.

**Supplement 9:**
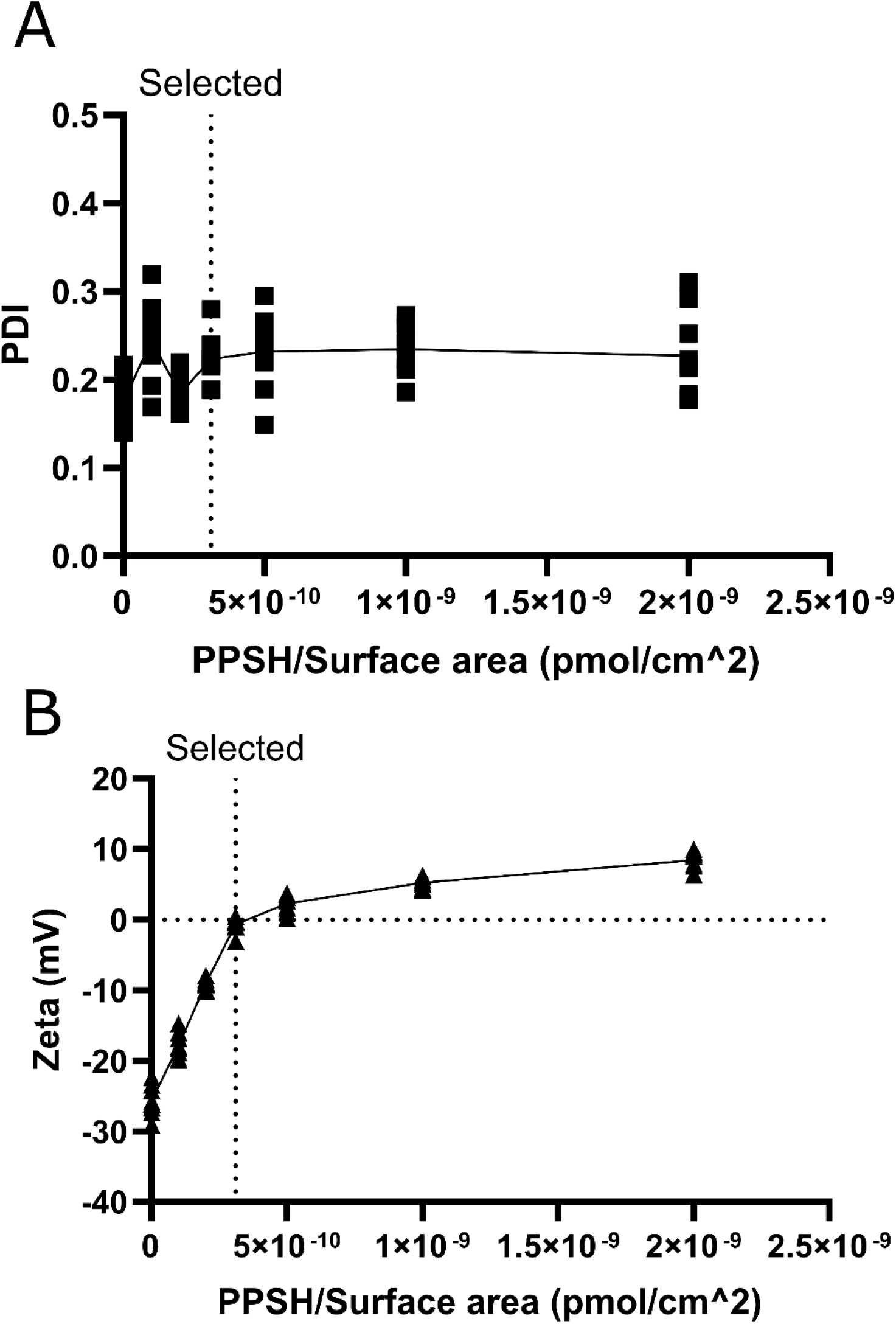
2^nd^ generation PPSH coating dose, PDI/Zeta potential. A) PDI for 16.5, 16.6 and 18.5 nm Au core sizes formulated at 23 pmol/cm^2 RNP then coated with a range of PPSH. B) Zeta potential for 2^nd^ Generation CRISPR-AuNP in 10 mM HEPES buffer at pH 7.4. For figures, n = minimum of 3, maximum of 12 technical replicates.

**Table 1:**
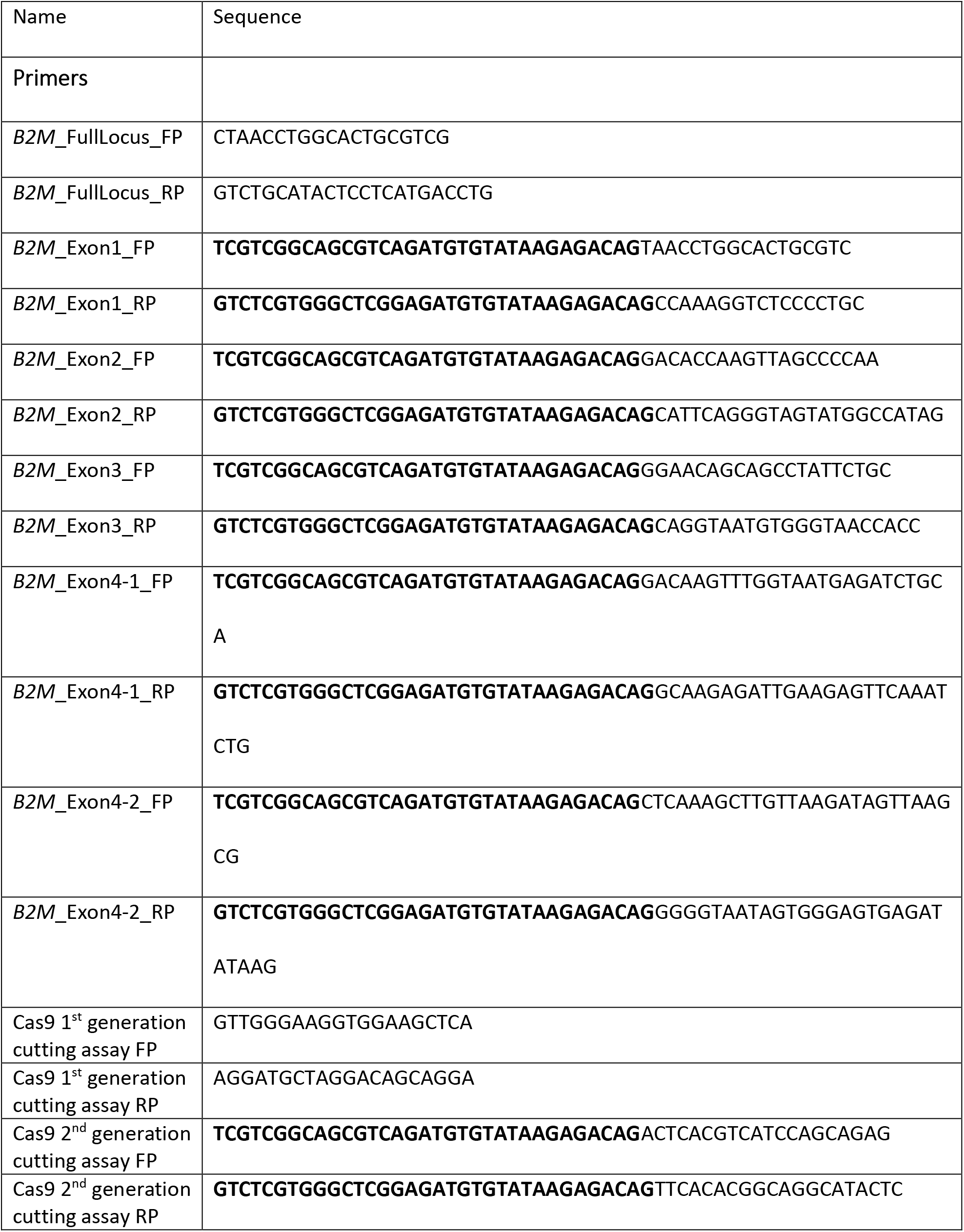
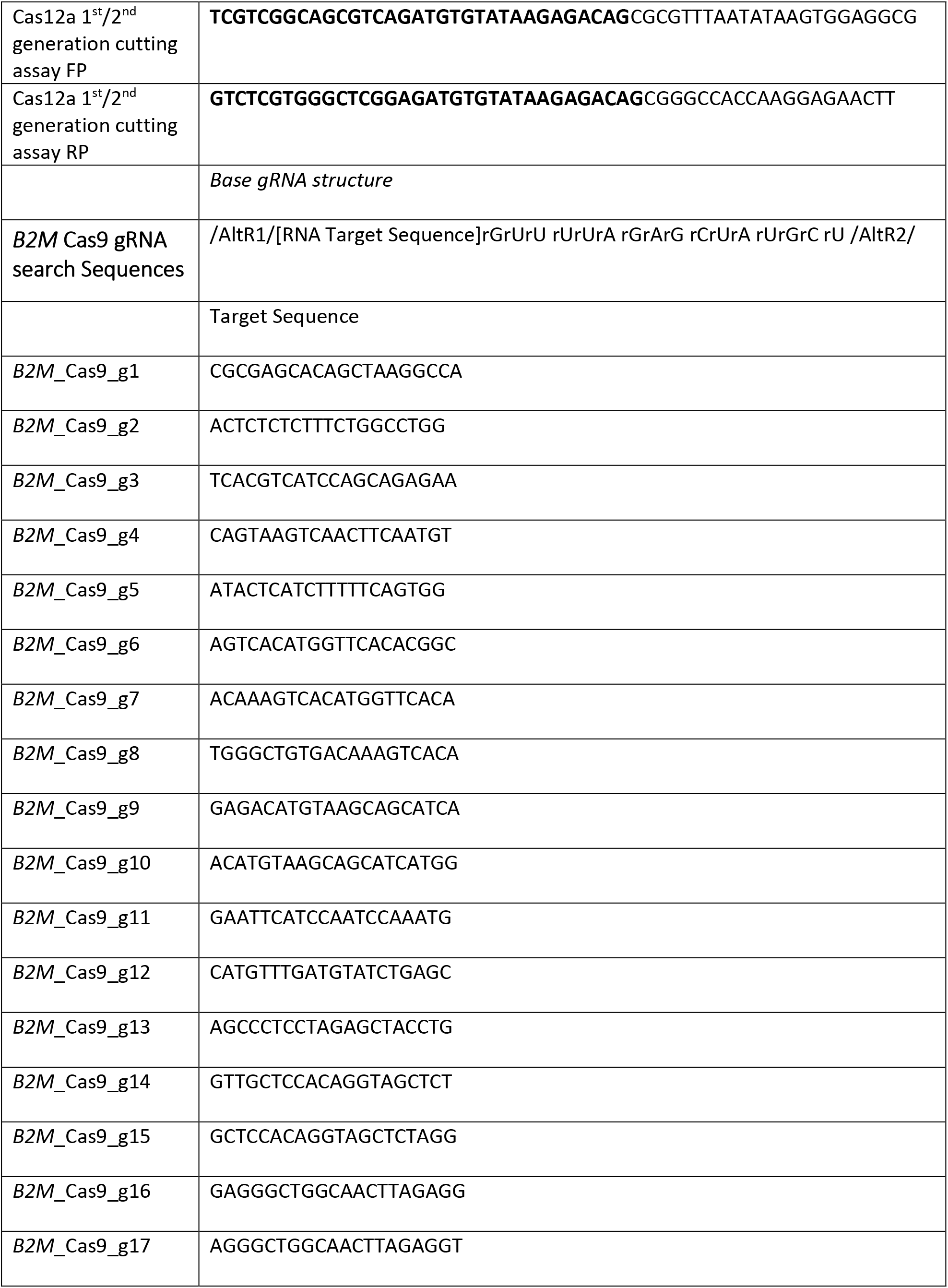
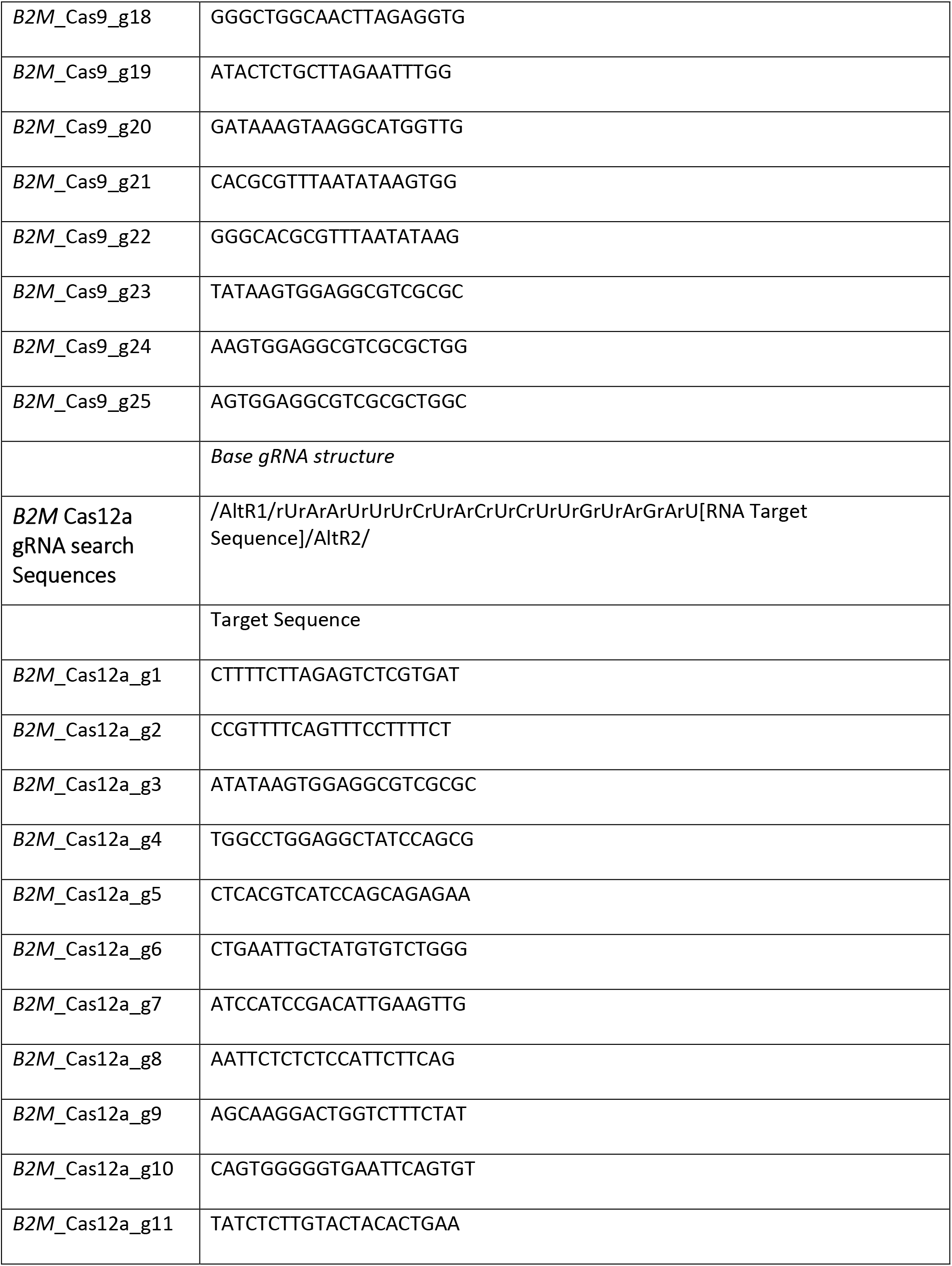
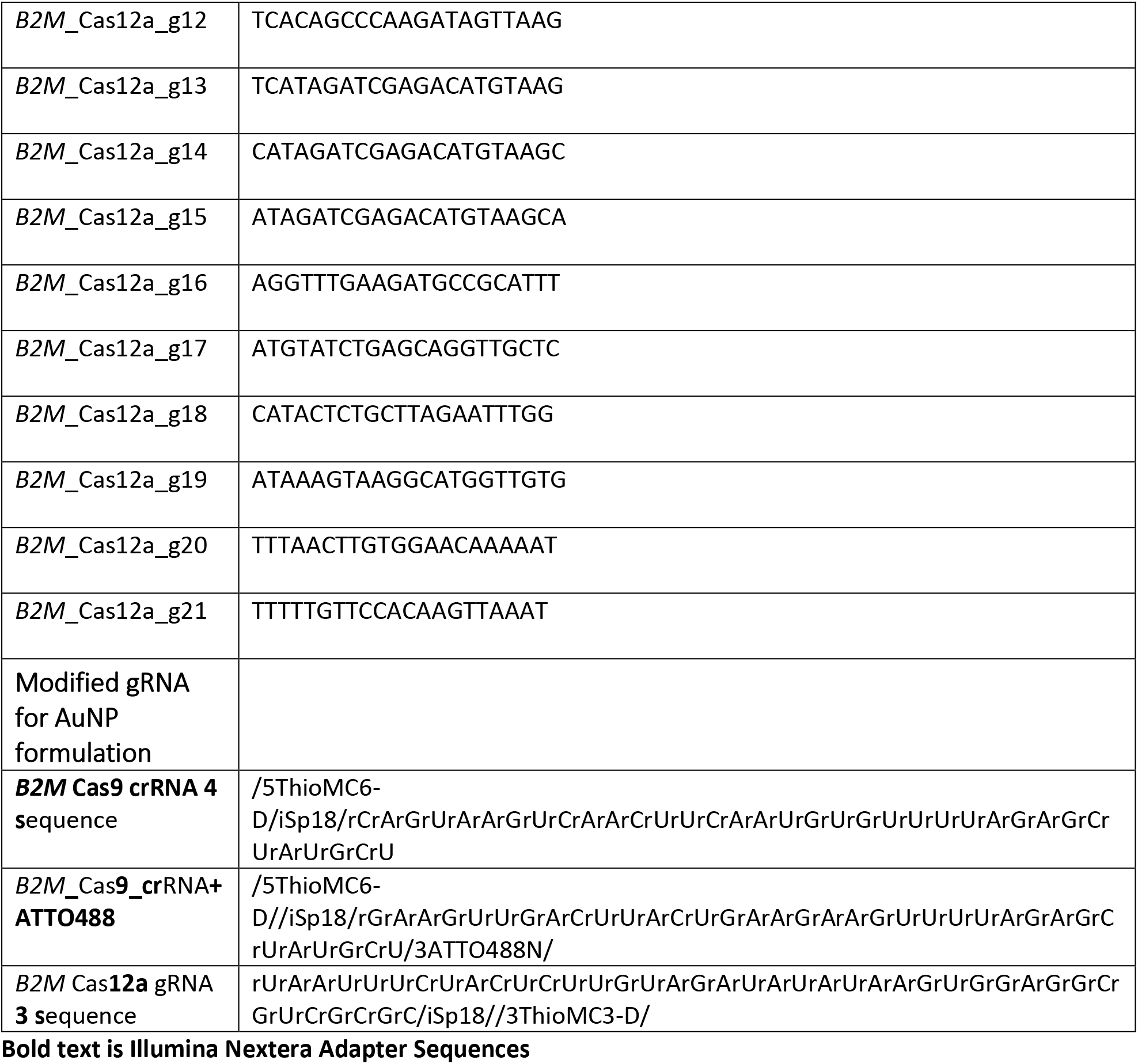
Oligonucleotide sequences.

**Table 2:**
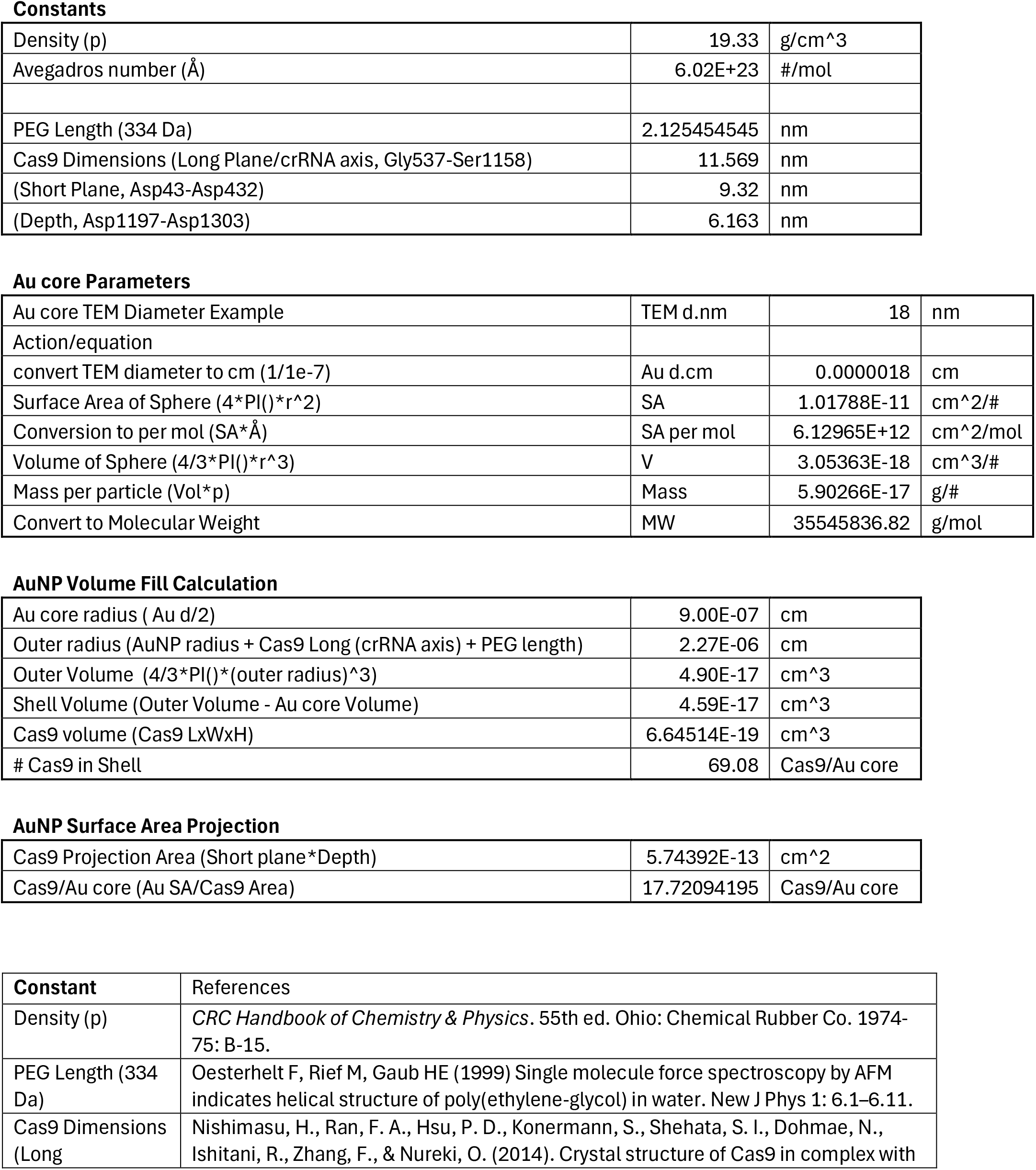
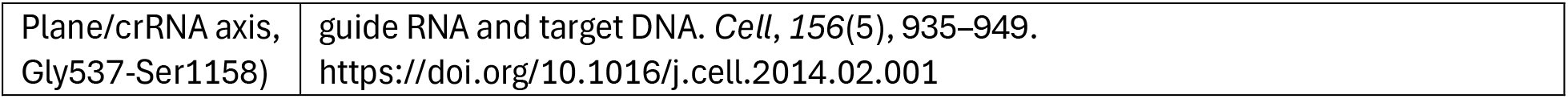
AuRNP Theoretical Coating Calculation.

