## Supplementary figures and images for "Cas9 RNP Physiochemical Analysis for Enhanced CRISPR-AuNP Assembly and Function"

### Supplement 1

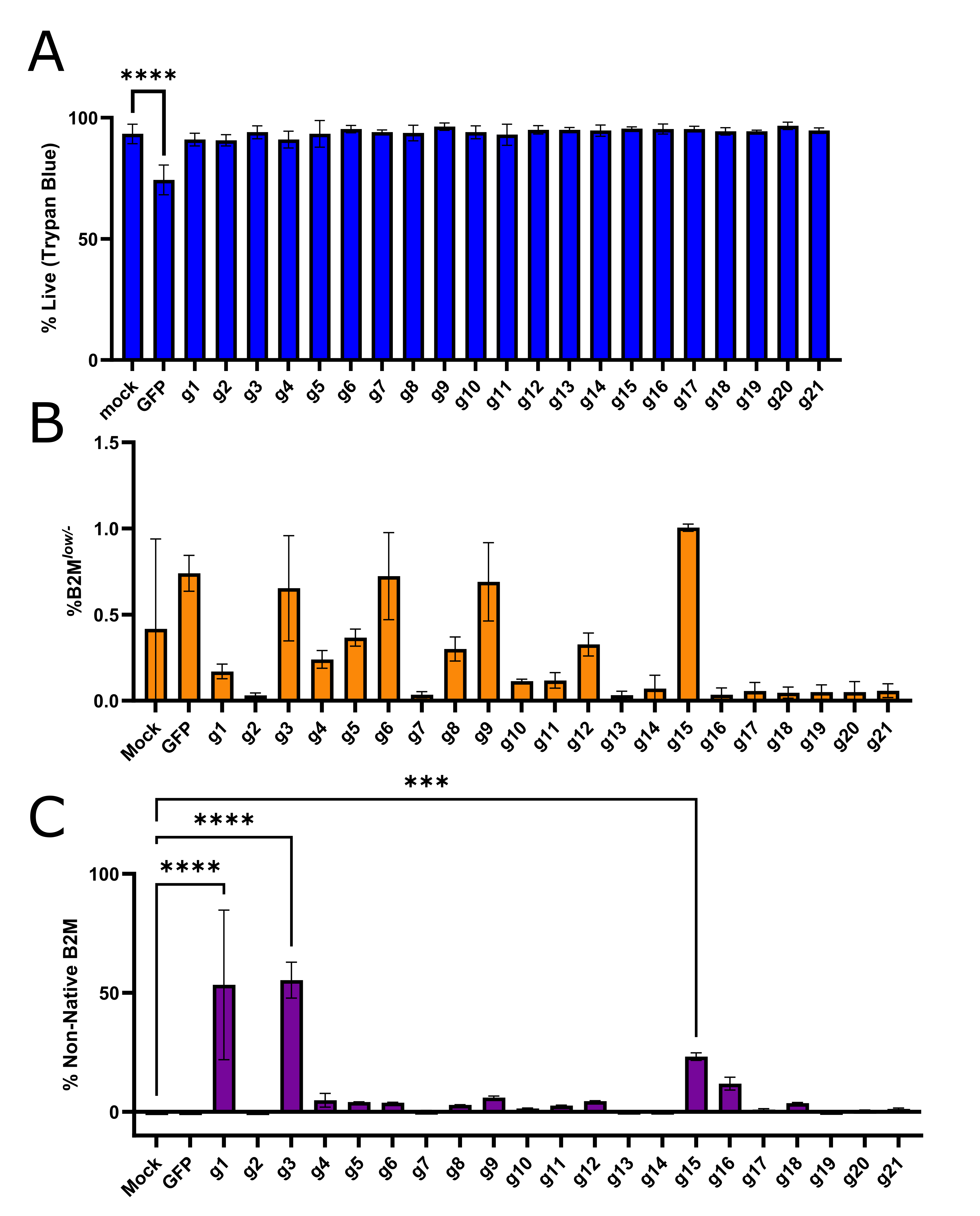

### Supplement 2

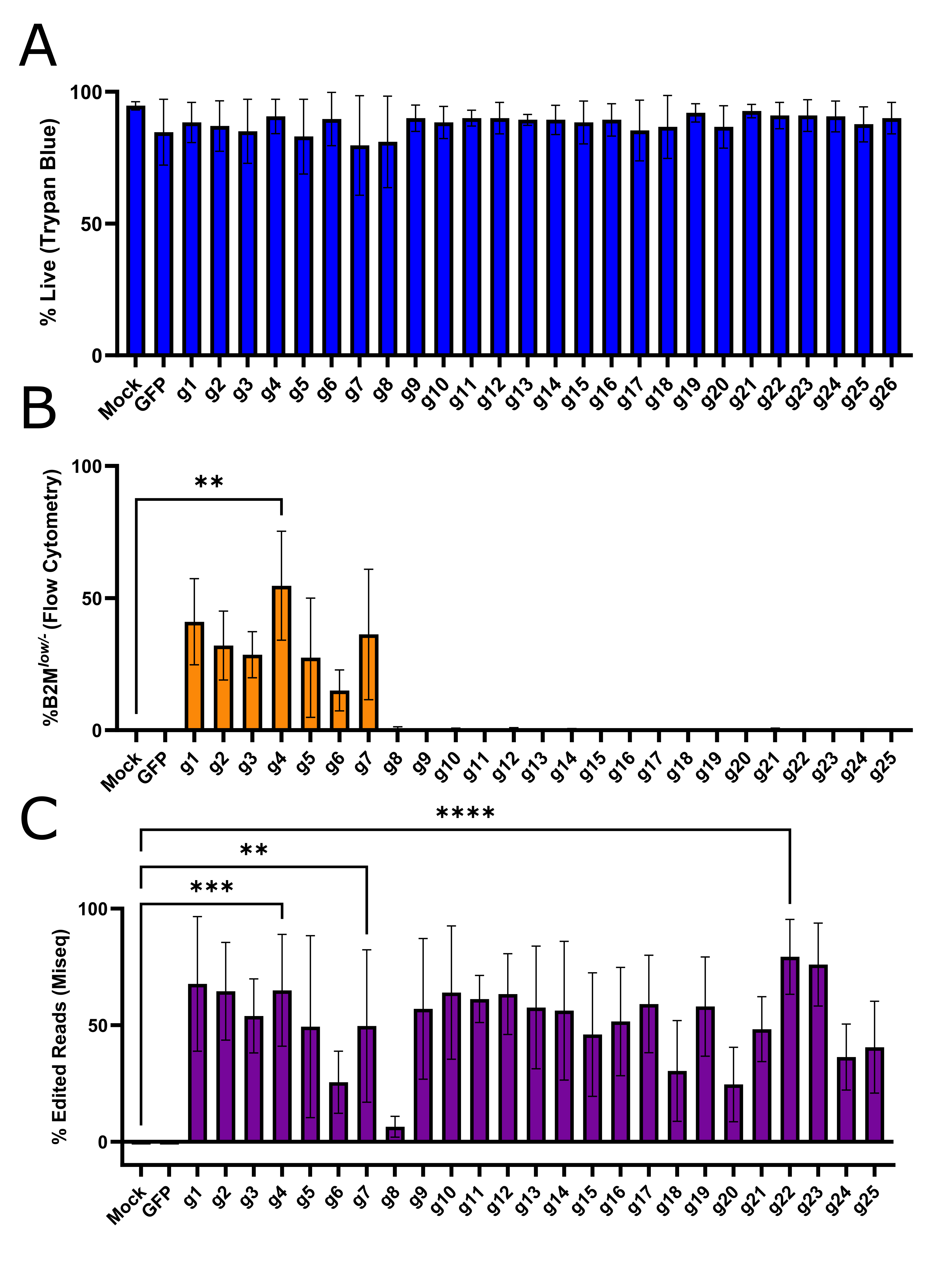

### Supplement 3

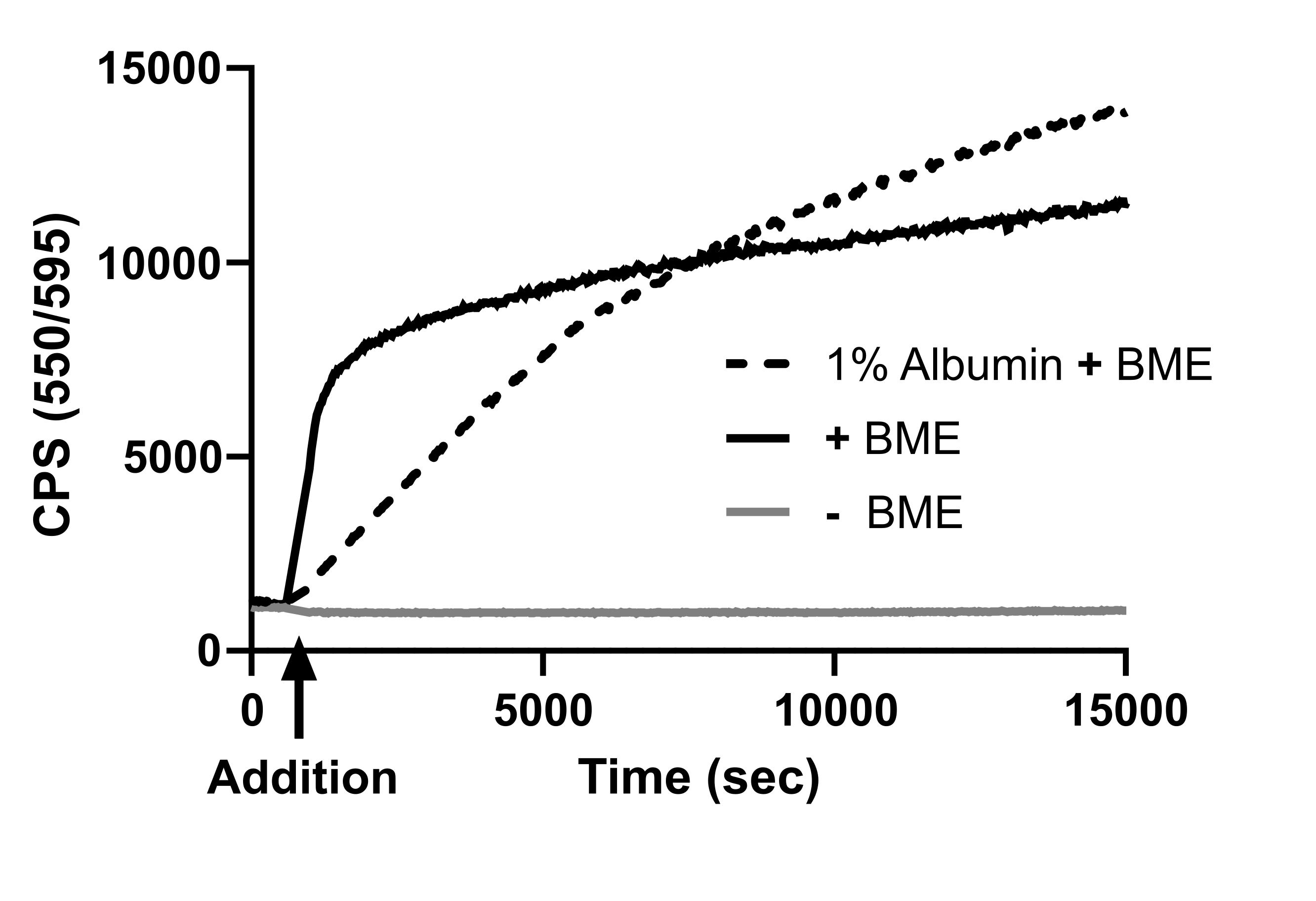

### Supplement 4

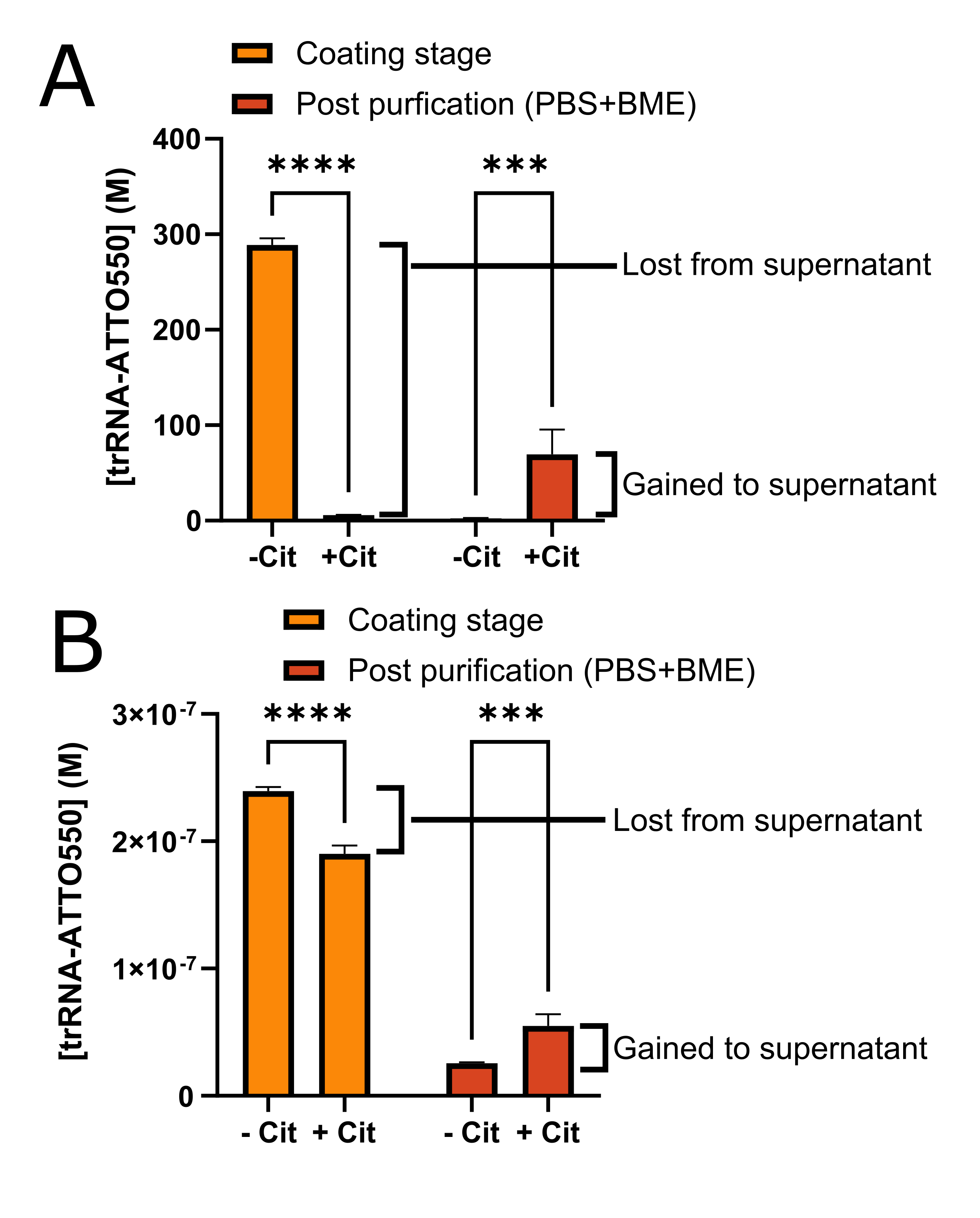

### Supplement 5

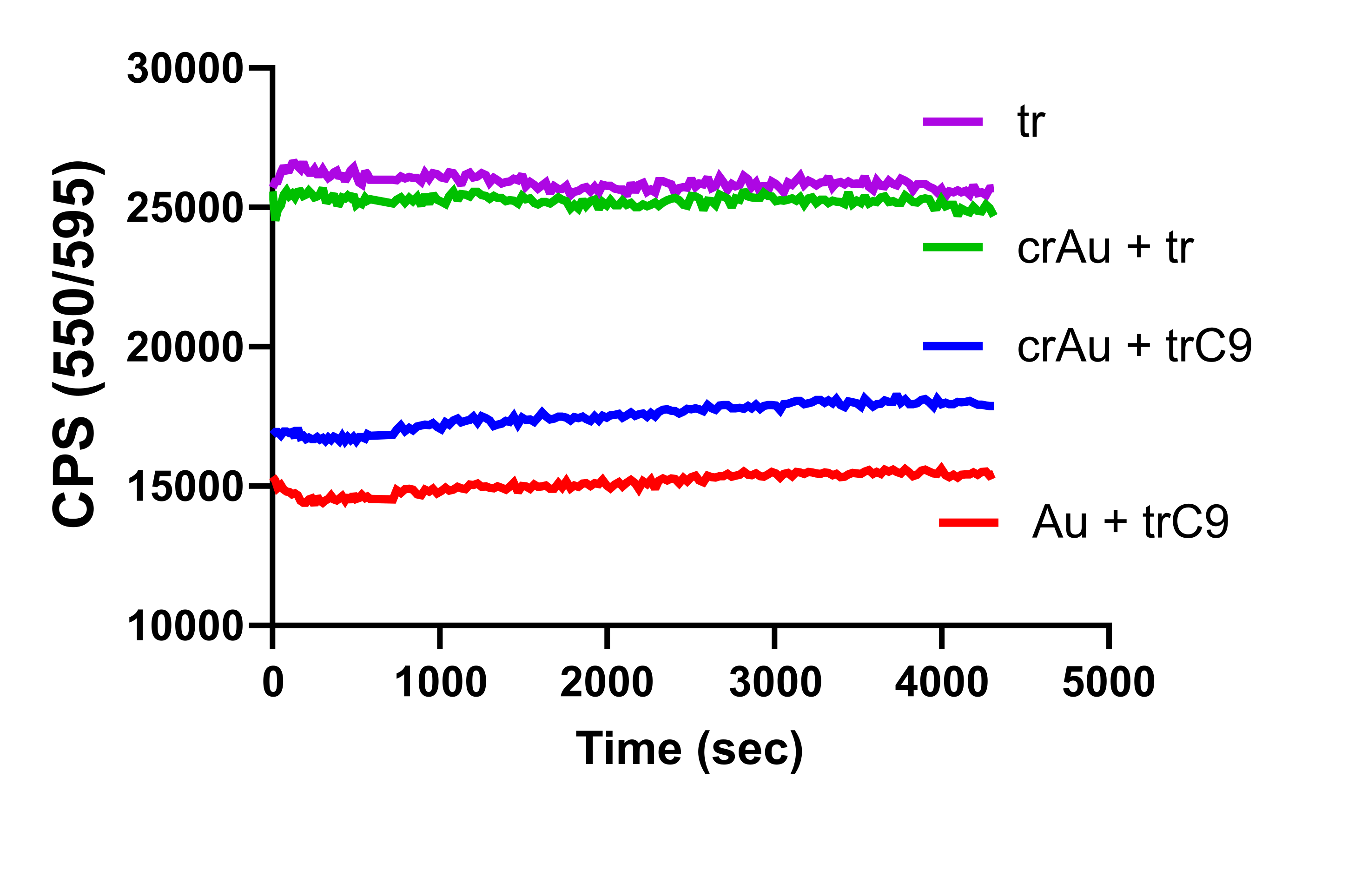

### Supplement 7

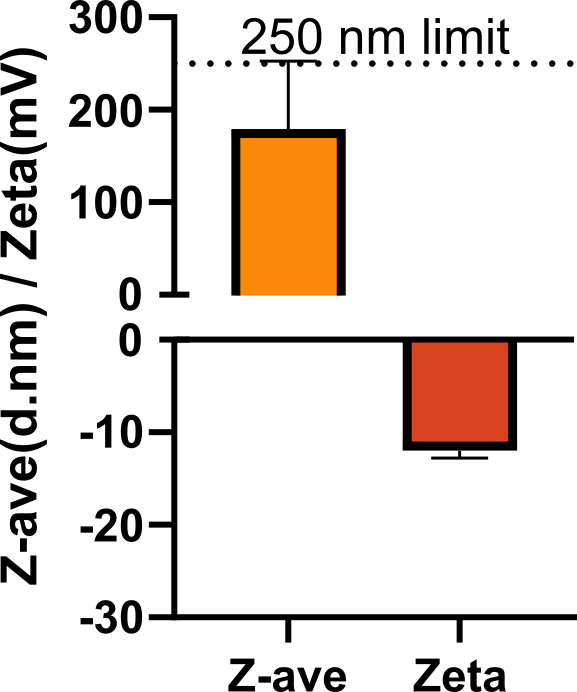

### Supplement 8

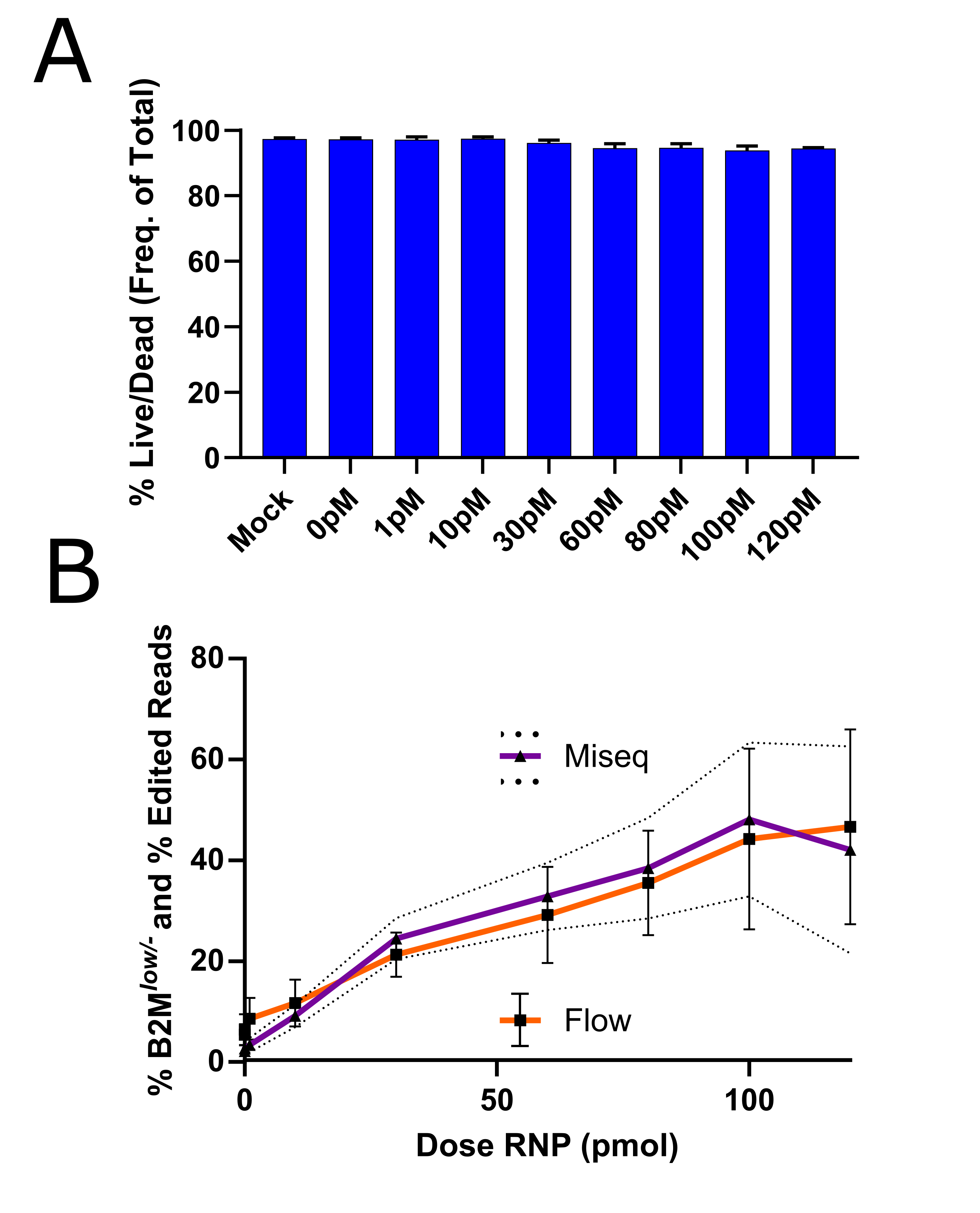

### Supplement 9

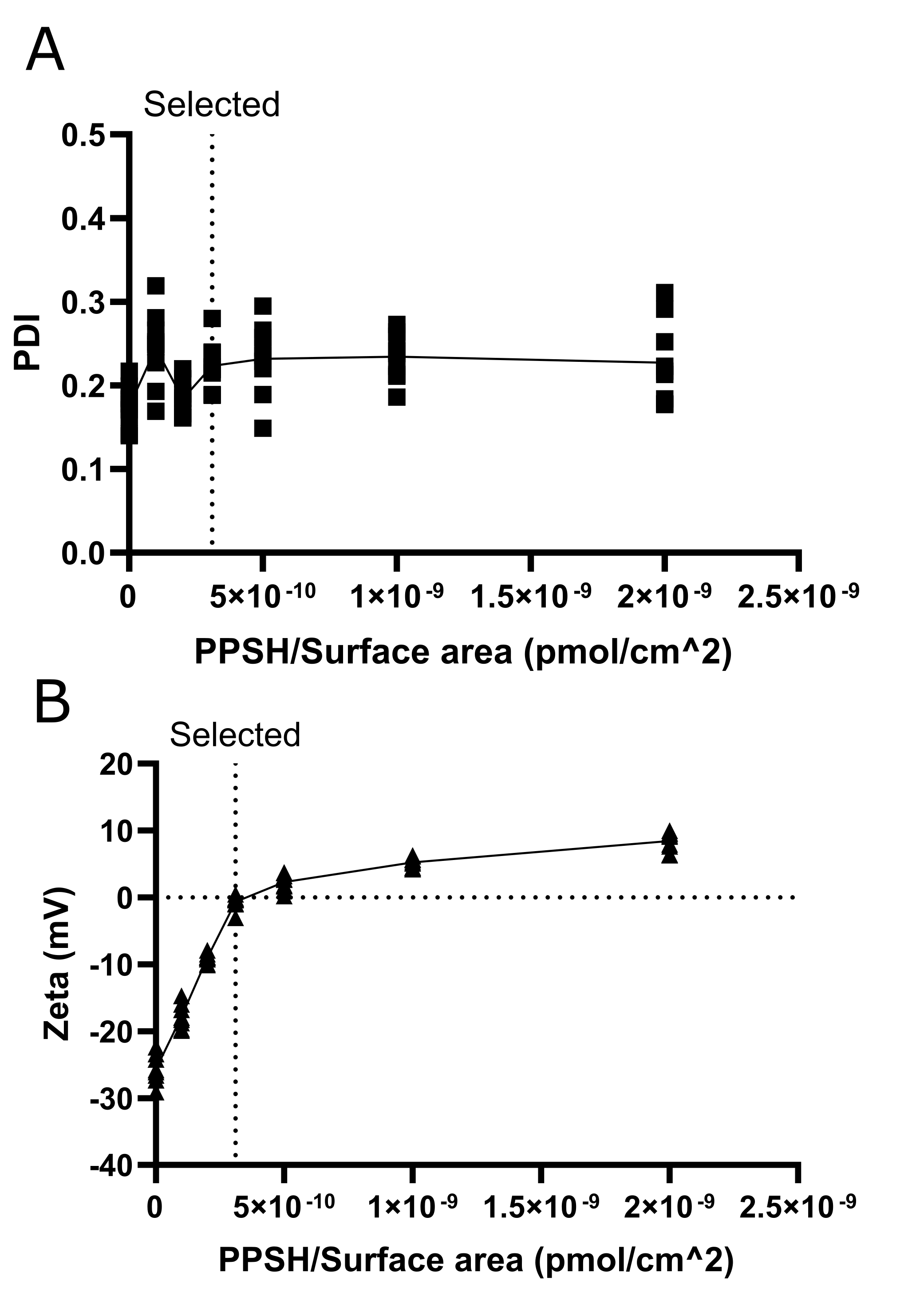
